# Visualization of the relative contributions of conductances in neuronal models with similar behavior and different conductance densities

**DOI:** 10.1101/427260

**Authors:** Leandro M. Alonso, Eve Marder

## Abstract

Conductance-based models of neural activity produce large amounts of data that can be hard to visualize and interpret. Here we introduce two novel visualization methods to display the dynamics of the ionic currents, and to investigate how the contribution of each current changes in response to perturbation. We explored the solutions of a single compartment, conductance-based model of neural activity with seven voltage-gated ionic currents and a leak channel. We employed landscape optimization to find sets of maximal conductances that produce similar target activity and displayed the dynamics of the currents. We examined in detail six examples of a bursting model neuron that differ as much as 3-fold in the conductance densities of each of the 8 currents in the model. The maximal conductance of each current does not simply predict the importance of the current for neuronal dynamics. We then compared the effects of systematically reducing the conductances of each current for neuronal dynamics, and demonstrate that models that appear similar under starting conditions behave dramatically differently to the decreases in conductance densities. These examples provide heuristic insight into why individuals with similar behavior can nonetheless respond widely differently to perturbations.

## I. Introduction

Experimental and computational studies have clearly demonstrated that neurons and circuits with very similar behaviors can nonetheless have very different values of the conductances that control intrinsic excitability and synaptic strength. Using a model of the crustacean stomatogastric ganglion (STG), Prinz et al. (2004) showed that similar network activity can arise from widely different sets of membrane and synaptic conductances. Recent experimental measurements have shown two to six-fold variability in individual components in the same identified neurons (Schulz et al., 2006, 2007; Roffman et al., 2011; Swensen and Bean, 2005). At the same time, the use of RNA sequencing and other molecular measurements have shown significant cell-to-cell variability in the expression of ion channels (Temporal et al., 2011, 2014; Tobin et al., 2009). Together these results suggests that these activities arise from different cellular and network mechanisms. Here we use conductance-based models to explore how different these mechanisms are and how they respond to perturbation.

Because of the intrinsic variability, canonical models that capture the mean behavior of a set of observations are not sufficient to address these issues (Golowasch et al., 2002; Balachandar and Prescott, 2018). In order to incorporate intrinsic biophysical variability Prinz et al. (2004) introduced an ensemble modeling approach. They constructed a database with millions of model parameter combinations, analyzed their solutions to assess network function, and screened for conductance values for which the activity resembled the data (Calabrese, 2018). One alternative to this approach was later introduced by Achard and De Schutter (2006). They combined evolutionary strategies with a fitness function based on a phase-plane analysis of the models’ solutions to find parameters that reproduce complex features in electrophysiological recordings of neuronal activity, and applied their procedure to obtain 20 very different computational models of cerebellar Purkinje cells. Here we adopt a similar approach and apply evolutionary techniques to optimize a different family of landscape functions that rely on thresholds or Poincaré sections to characterize the models’ solutions.

In some respects, biological systems are a black-box because one cannot read out the values over time of all the underlying components. In contrast, computational models allow us to inspect how all the components interact and this can be used to develop intuitions and predictions about how these systems will respond to extreme perturbations. Despite this, much modeling work focuses on the variables of the models that are routinely measured in experiments, such as the membrane potential. While in the models we have access to all state variables, this information can be hard to represent when many conductances are at play. Similarly, the effect of perturbations – such as the effect of partially or completely removing a particular channel – can be complex and also hard to display in an compact fashion. Here we address these difficulties and propose two novel visualization methods. We represent the currents in a model neuron using stacked area plots: at each time step we display the shared contribution of each current to the total current through the membrane. This representation is useful to visualize which currents are most important at each instant and allows the development of insight into how these currents behave when the system is perturbed. Perturbation typically results in drastic changes of the waveform of the activity and these changes depend on the kind of perturbation under consideration. We developed a representation that relies on computing the classical probability of *V* (*t*), which allows the visualization of these changes. We illustrate the utility of these procedures using models of single neuron bursters or oscillators.

## II. Results

### Finding parameters: landscape optimization

The numerical exploration of conductance-based models of neurons is a commonplace approach to address fundamental questions in neuroscience (Dayan and Abbott, 2001). These models can display much of the phenomenology exhibited by intracellular recordings of single neurons and have the major advantage that many of their parameters correspond to measurable quantities (Herz et al., 2006). However, finding parameters for these models so that their solutions resemble experimental observations is a difficult task. This difficulty arises because the models are nonlinear, they have many state variables and they contain a large number of parameters (Bhalla and Bower, 1993). These models are complex and we are not aware of a general procedure that would allow the prediction of how an arbitrary perturbation in any of the parameters will affect their solutions. The problem of finding sets of parameters so that a nonlinear system will display a target behavior is ubiquitous in the natural sciences. A general approach to this problem consists of optimizing a score function that compares features of the models’ solutions to a set of target features. Consequently, landscape-based optimization techniques for finding parameters in compartmental models of neurons have been proposed before (Achard and De Schutter, 2006; Druckmann et al., 2007; Ben-Shalom et al., 2012). Here we employ these ideas to develop a family of score functions that are useful to find parameters so that their activities reach a desired target.

In this work we started with a well-studied model of neural activity described previously (Liu et al., 1998; Goldman et al., 2001; Prinz et al., 2004; OLeary et al., 2014). The neuron is modeled according to the Hodgkin-Huxley formalism using a single compartment with eight currents. Following Liu et al. (1998), the neuron has a sodium current, *I_Na_;* transient and slow calcium currents, *I_CaT_* and *Ic_a_s*; a transient potassium current, *I_A_;* a calcium-dependent potassium current, *I_KCa_*; a delayed rectifier potassium current, *I_Kd_*; a hyperpolarization-activated inward current, *I_H_*; and a leak current *I_leak_*.

We explored the space of solutions of the model using landscape optimization. The procedure consists of three parts. First we generate voltage traces by integration of eq. (5) (methods). We then score the traces using an objective function that defines a target activity. Finally, we attempt to find minima of the objective function. The procedures used to build objective functions whose minima correspond to sets of conductances which yield the target activities are shown in Figure 1. Voltage traces were generated by integration of eq. (5) from fixed initial conditions (supplementary) and were then scored according to a set of simple measures. The procedure is efficient in part because we chose measures that require little computing power and yet are sufficient to build successful target functions. For example, we avoid the use of Spike Density Functions (SDF) and Fourier transforms when estimating burst frequencies and burst durations. In this section we describe target functions whose minima correspond to bursting and tonic activity in single compartment models. Our approach can also be applied to the case of small circuits of neurons (Prinz et al., 2004).

**FIG. 1.**
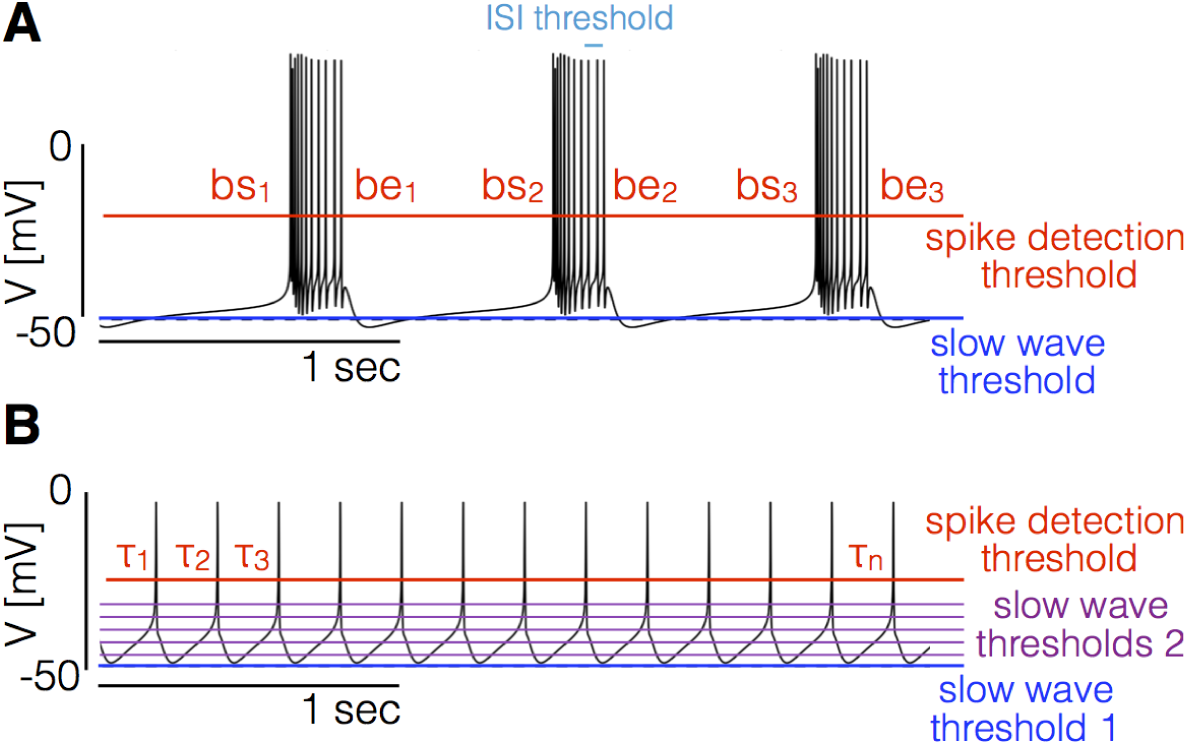
Landscape optimization can be used to find models with specific sets of features. **A** Example model bursting neuron. The activity is described by the burst frequency and the burst duration in units of the period (duty cycle). The spikes detection threshold (red line) is used to determine the spike times. The ISI threshold (cyan) is used to determine which spikes are bursts starts (bs) and bursts ends (be). The slow wave threshold (blue line) is used to ensure that slow wave activity is separated from spiking activity. **B** Example model spiking neuron. We use thresholds as before to measure the frequency and the duty cycle of the cell. The additional slow wave thresholds (purple) are used to control the waveform during spike repolarization.

We begin with the case of bursters (Fig. 1A). We targeted this type of activity by measuring the bursting frequency, the duty cycle, and the number of crossings at a threshold value to ensure that spiking activity is well separated from slow wave activity. To measure the burst frequency and duty cycle of a solution we first compute the time stamps at which the cell spikes. Given the sequence of values *V* = {*V_n_*} we determine that a spike occurs every time that *V* crosses the spike detection threshold *T_sp_* = −20mV (red in Fig 1). We build a sequence of spike times *S* = {*s_i_*} by going through the sequence of voltages {*V_n_*} and keeping the values of *n* for which *V_n_ ≤ T_sp_* and *V_n+1_ > T_sp_* (we consider upward crossings). Each element *s_i_* of the sequence *S* contains the time step at which the i-th spike is detected. Bursts are determined from the sequence of spike times *S*; if two spikes happen within a temporal interval shorter than *δ_spt_* = 100*msec* they are part of a burst. Using this criteria we can find which of the spike times in *S* correspond to the start and end of bursts. The starts (bs) and ends (be) of bursts are used to estimate the duty cycle and burst frequency. We loop over the sequence of spike times and determine that a burst start at *s_i_* if *s_i+_*_1_ − *s_i_* < *δ_spt_* and *s_i_* − *s_i_*_−1_ > *δ_spt_*. After a burst starts, we define the end of the burst at *s_k_* if *s_k_*_+1_ − *s_k_ > δ_spt_* and *s_k_ − s_k−_*_1_ < *δ_spt_*. When a burst ends we can measure the burst duration as *δ_b_* = *s_k_ − s_i_*, and since the next burst starts (by definition) at *s_k_*_+1_ we also can measure the “period” (if periodic) of the oscillation as *τ_b_* = *δ_b_* + (*s_k+_*_1_ − *s_k_*). Every time a burst starts and ends we get an instance of the burst frequency 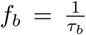 and the duty cycle 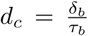. We build distributions of these quantities by looping over the sequence *S* and define the burst frequency and duty cycle as the mean values *< f_b_ >* and < *dc* >. Finally, we count downward crossings in the sequence *V_n_* with *two* slow wave thresholds *#_sw_* (with *t_sw_* = −50 ± 1*mV*) and the total number of bursts #_b_ in *S*.

For any given set of conductances we simulated the model for 20 seconds and dropped the first 10 seconds to mitigate the effects of transient activity. We then computed the burst frequency *< f_b_* >, the duty cycle *< dc >*, the number of crossings with the slow wave thresholds *#_sw_* and the number of bursts #*_b_*. We discard unstable solutions; a solution is discarded if *std*({*fb*}) *≥< f_b_ >* ×0.1 or *std*({*dc*}) ≥< *dc* > ×0.2. If a solution is not discarded we can use the following quantities to measure how close it is to the target behavior,

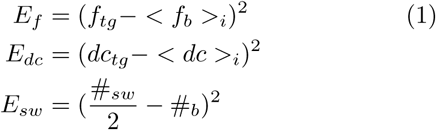

Here *E_f_* measures the mismatch of the bursting frequency of the model cell with a target frequency *f_tg_* and *E_dc_* accounts for the duty cycle. *E_sw_* measures the difference between the number of bursts and the number of crossings with the slow wave thresholds *t_sw_* = −50 ± 1*mV*; because we want a clear separation between slow wave activity and spiking activity, we ask that *#_sw_* = #*_b_*. Note that if during a burst *V* goes below *t_sw_* this solution would be penalized (factor 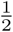 accounts for using two slow wave thresholds). Let ***g*** denote a set of parameters, we can then define an objective function

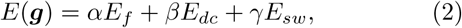

where the weights (*α, β, γ*) determine the relative importance of the different sources of penalties. In this work we used *α* =10, *β* = 1000, *γ* = 10, and the penalties *E_i_* were calculated using *T* =10 seconds with *dt* = 0.1 msecs. The target behavior for bursters was defined by *dc_tg_* = 0.2 (duty cycle 20%) *(dc_tg_* = 0.2) and bursting frequency *f_tg_* = *1Hz*.

We can use similar procedures to target tonic spiking activity. Note that the procedure we described previously to determine bursts from the sequence of spike times *S* is useful in this case too. If a given spike satisfies our definition of burst start and it also satisfies the definition of burst end then it is a single spike and the burst duration is zero. Therefore we compute the bursts and duty cycles as before and ask that the the target duty cycle is zero.

There are multiple ways to produce tonic spiking in this model and some solutions display very different slow wave activity. To further restrict the models we placed a middle threshold at *t_mid_* = −35*mV* and detected downward crossings at this value. We defined *E_lag_* as the lag between the upward crossings at the spiking threshold (*t_spk_* = − 20*mV*) and downward crossings at *t_mid_. E_lag_* is useful because it takes different values for tonic spikers than it does for single-spike bursters even though their spiking patterns can be identical. Finally, we found that the model attempts to minimize *E_lag_* at the expense of hyperpolarizing the membrane beyond − 50*mV* and introducing a wiggle that can be different in different solutions. In order to penalize this we included additional thresholds between −35*mV* and −45*mV*, counted the number downward crossings at these values *#_midi_*, and asked that these numbers are equal to the number of spikes #_s_. With these definitions we define the partial errors as before,

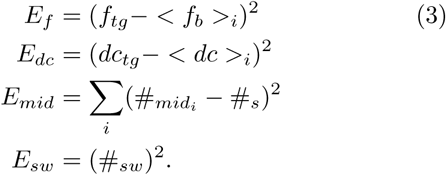

The total error as a function of the conductances reads as follows,

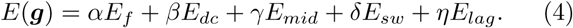

The values *α* = 1000, *β* = 1000, *γ* = 100, *δ* = 100 and *η* = 1, produce solutions that are almost identical to the one displayed in Fig. (1B).

In all cases, evaluation of the objective functions requires that the models are simulated for a number of seconds and this is the part of the procedure that requires most computing power. Longer simulations will provide better estimations for the burst frequency and duty cycle of the cells, but will linearly increase the time it takes to evaluate the objective function. If the simulations are shorter, evaluations of the objective function are faster but its minimization may be more difficult due to transient behaviors and its minima may not correspond to stable solutions. In this work we minimized the objective function using a custom genetic algorithm (Holland, 1992; Goldberg and Holland, 1988). The choice of the optimization routine and the choice of the numerical scheme for the simulations are independent of the functions. The same functions can be utilized to estimate parameters in models with different channel types.

### Visualizing the dynamics of ionic currents: currentscapes

Most modeling work focuses on the variables of the models that are routinely measured in experiments such as the membrane potential. While in the models we have access to all state variables, this information can be hard to represent when several current types are at play. One difficulty is that some currents like *Na* and *Kd* vary over several orders of magnitude, while other currents like the *leak* and *H* span smaller ranges. Additionally, the relative contribution of each current to the total flux through the membrane is different at different times. Here we introduce a novel representation that is simple and permits displaying the dynamics of the currents in a cohesive fashion.

At any given time stamp we can compute the total inward and outward currents. We can then express the values of each current as a percentage of this quantity. The normalized values of the currents at any time can be displayed as a pie chart representing the share of each current type. Because we want to observe how these percentages change in time, we display the shares in a bar instead of a disk. The currentscapes are constructed by applying this procedure to all time stamps and stacking the bars. These types of plots are known as stacked area plots and their application to this problem is novel. Figure 2 shows the currentscape of a periodically bursting model neuron over one cycle.

**FIG. 2.**
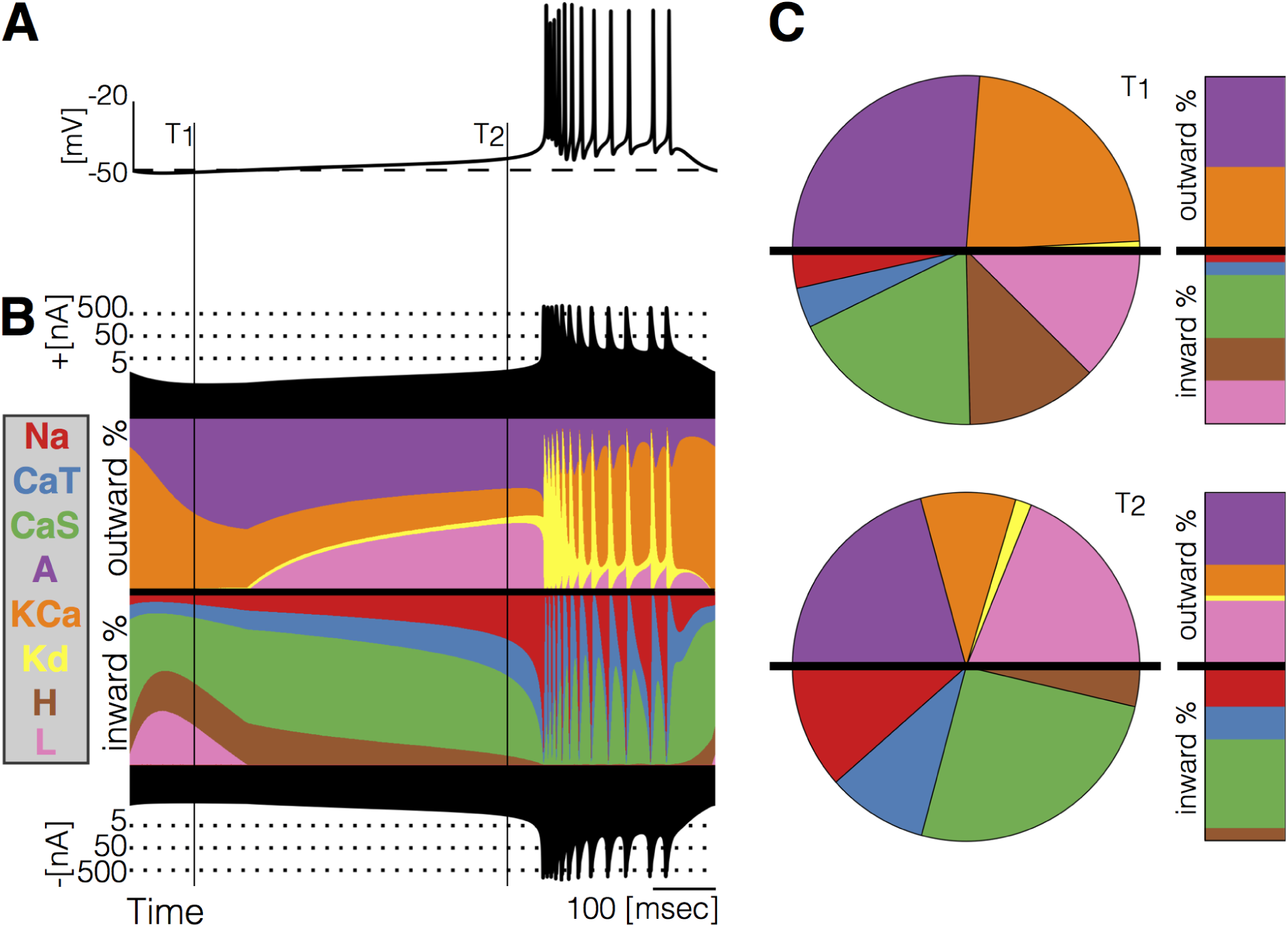
Currentscape of a model bursting neuron. **A** simple visualization of the dynamics of ionic currents in these models A Membrane potential of a periodic burster. **B** Percent contribution of each current to the total outward and inward currents at each time stamp. The black filled curves on the top and bottom indicate total inward outward currents respectively on a logarithmic scale. The color curves show the time evolution of each current as a percentage of the total current at that time. For example, at *t* = *T*_1_ the total outward current is ≈ 2.5*nA* and the orange shows a large contribution of *KCa*. At *t* = *T*_2_ the total outward current has increased to ≈ 4*nA* and the *KCa* current is contributing less to the total. **C** Percent contribution of each current type to the total inward and outward currents displayed as pie charts at times *T*_1_ and *T*_2_

### Visualizing changes in the waveforms as a parameter is changed

To visualize changes in the activity as a conductance is gradually removed we computed the distribution of membrane potential *V* values. This reduction contains information about the waveform of the membrane potential, while all temporal information such as frequency can no longer be recovered. The number of times that a given value of *V* is sampled is proportional to the time the system spends at that value. The distribution of *V*, shown in Fig. 3A, is larger than 10^4^ for values between −52*mV* and −40*mV*, and smaller than 10^3^ for *V* between −35*mv* and 20*mV*. The areas of the shaded regions are proportional to the probability that the system will be observed at the corresponding *V* range. Note that the area of the dark gray region is 10^5^ while the light gray is 0.5 × 10^4^, so the probability that the cell is, at any given time, in a hyperpolarized state is more than 20 times larger than the probability that the cell is spiking. The distribution shows the overall or total amplitude of the oscillation since the count of *V* is zero outside a range (−52*mV* to 20*mV*). As conductances are gradually removed the waveform of the activity changes and so does the distribution of *V* values. In Figure 3 we display the changes in these distributions as *gNa* is decreased.

**FIG. 3.**
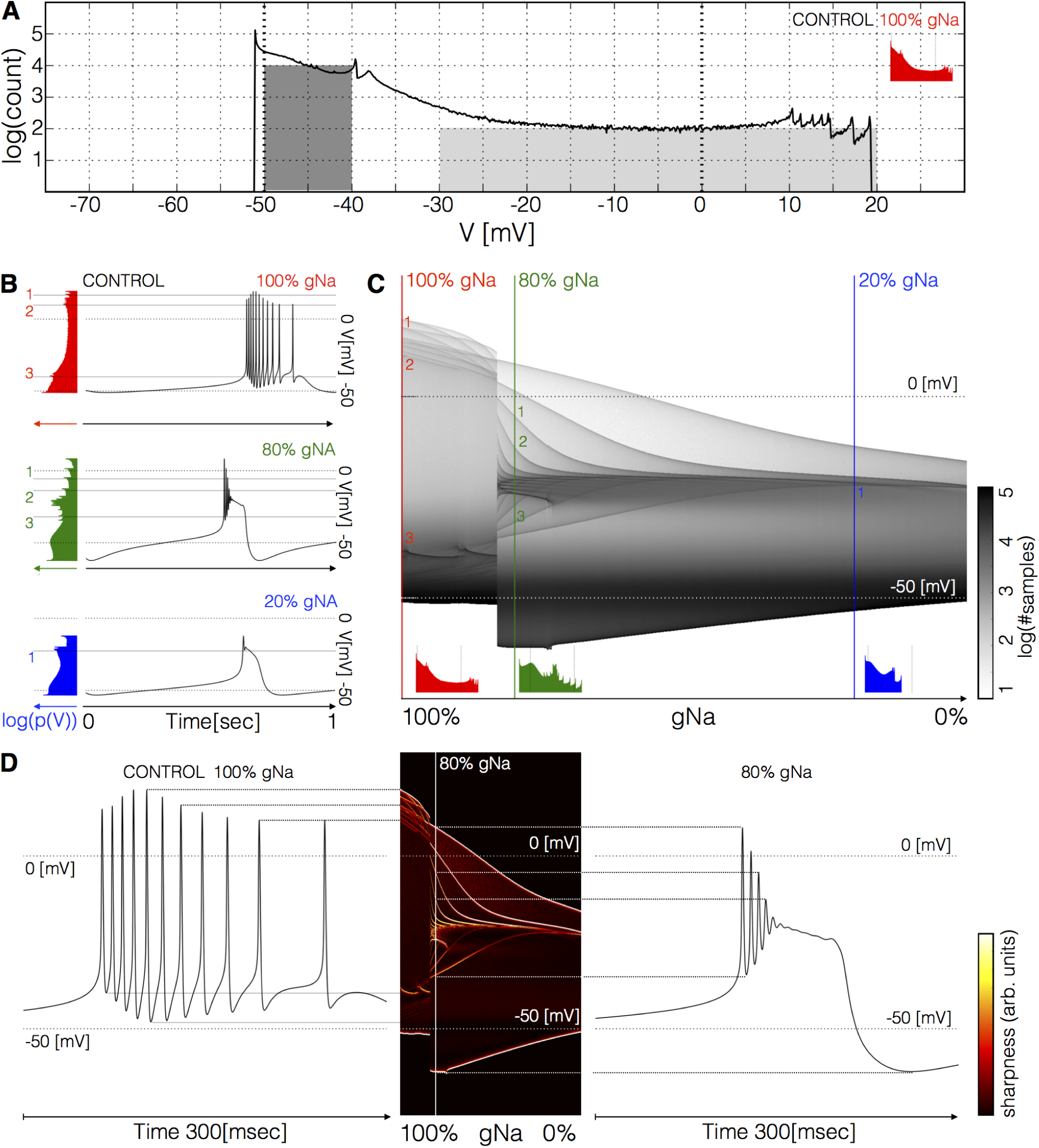
Membrane potential *V* distributions. **A** Distribution of membrane potential *V* values. The total number of samples is *N =* 2.2 × 10^9^. Y-axis is logarithmic. The area of the dark shaded region can be used to estimate of the probability that the activity is sampled between −50*mV* and −40*mV*, and the area of the light shaded region is proportional to the probability that *V*(*t*) is sampled between −30*mV* and 20*mV*. The area of the dark region is 20 times larger than the light region. **B** Membrane potential *V* and distribution of *V* (in colors) for three values of the parameter. The gray lines highlight that the amplitudes of the spikes produce peaks in the distributions. The system is sampled more often at values of *V* where it spends more time; when *V*(*t*) reaches a maximum or a minimum the velocity 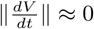 is small so it is more likely to obtain a sample at these values. **C** Distribution of *V* as a function of *V* and parameter *gNa*. The color lines indicate the values of *gNa* that correspond to the traces and distributions shown in **B. D** Waveforms under two conditions and their correspondence to the ridges of the distribution of *V*.

The distribution of *V* features sharp peaks. In many cases, the peaks in these distributions correspond to features of the waveform, such as the amplitudes of the individual spikes, or the minimum membrane potential (see Fig, 3B). This happens because every time the membrane potential reaches a maxima or minima (in time) the derivative 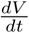 is close to zero. The system spends more time close to values of *V* where the velocity 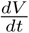 is small than in regions where 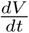 is large, as it occurs during the flanks of spikes. Therefore, when we sample a solution at a random instant, it is more likely that *V* corresponds to the peak of a spike than to either flank of the spike, while the most likely outcome is that *V* is in the hyperpolarized range (< −40*mV*). Notice also that in this particular burster there are 12 spikes in the burst but there are only 7 peaks in the distribution (between 10*mV* and 20*mV*); some spikes have similar amplitudes so they add to a larger peak in the distribution. This is not the case for the 80% condition where each peak corresponds to a maxima or minima of different amplitudes. Figure 3C shows the distributions of *V* as *gNa* is decreased. For each value in the range (1 to 0 with *N* = 1001 values) we computed the count *p*(*V*,*gNa*). The plot shows the distribution *log*_10_(*p*(*V*, *gNa*) + 1) in gray scales. In this example, the cell remains in a bursting regime up to ≈ 80%. The spikes produce thin ridges that show how their individual amplitudes change. Notice the abrupt change in membrane potential as the activity switches from bursting (control) to a single-spike bursting mode (80%*gNa*). After the transition the amplitude of the spikes are different; two spikes go beyond 0*mV* and the rest accumulate near −25*mV*. As *gNa →* 0 the oscillations collapse onto a small band at ≈ −20*mV* and only one spike is left.

The distributions allow the visualization of the amplitudes of the individual spikes, the slow waves, and other features as the parameter *gNa* is changed. To highlight ridges in the distributions, the center panel in Figure 3D show the derivative *∂_v_log*_10_(*p*(*V*)) in colors. This operation is similar to performing a Sobel filtering(Sobel and Feldman, 1968) of the image in Fig. 3C. The traces on each side of this panel correspond to the control (right) and *80%gNa* conditions. Notice how the amplitudes of each spike, features of the slow wave, and overall amplitude correspond to features in the probability distribution. This representation permits displaying how the features of the waveform change for many values of *gNa*.

### The maximal conductances do not fully predict the currentscapes

We explored the solutions of a classic conductance-based model of neural activity using landscape optimization and found many sets of parameters that produce similar bursting activity. Inspired by intracellular recording performed in *PD* neurons in crabs and lobsters we targeted bursters with frequencies *f_b_* ≈ 1*Hz* and duty cycles *dc* ≈ 20%. We built *N* = 1000 bursting model neurons and inspected the dynamics of their currents using their currentscapes. Based on this, we selected six models that display similar membrane activity via different current compositions for further study. Because the models are nonlinear, the relationship between the dynamics of a given current type and the value of its maximal conductance is non-trivial. Figure 4 shows the values of the maximal conductances in the models (top) and their corresponding activity together with their currentscapes (bottom).

**FIG. 4.**
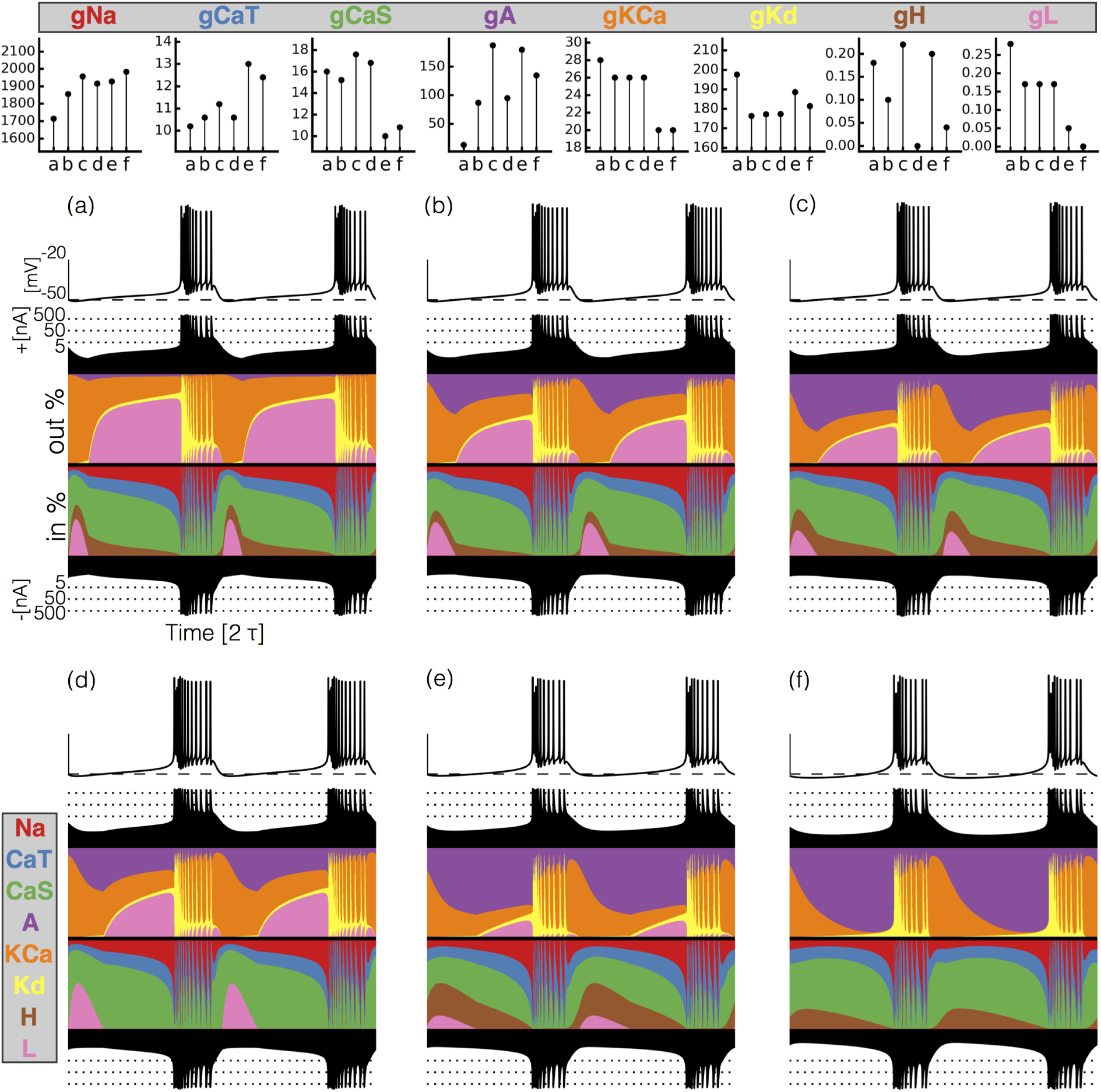
Currentscapes of model bursting neurons. (top) Maximal conductances of all model bursters. (bottom) The panels shows the membrane potential of the cell and the percent contribution of each current over two cycles.

It can difficult to predict the currentscapes based on the values of the maximal conductances. In most cases it appears that the larger the value of the maximal conductance, the larger the contribution of the corresponding current. However, this does not hold for some current types in these models. For example, burster (f) shows the largest *A* contribution, but bursters (c) and (e) have larger values of *gA*. The maximal conductance of the *CaS* current is low in model (f) but the contribution of this current to the total is similar to that in models (a) and (b). The values of *gKCa* are similar for bursters (e) and (f) but the dynamics of this current is visibly different.

### Response to current injection

The models produce similar activity with different current dynamics. To further expose differences in how these activities are generated we subjected the models to simple perturbation paradigms. We begin describing the response to constant current injections in Figure 5. Figures 5A and 5B show the membrane potential of model (a) for different values of injected current. In control, the activity corresponds to regular bursting and larger depolarizing currents result in a plethora of different regimes. The distributions of inter-spike intervals (ISI) provide a mean to characterize these regimes (Fig. 5C). When the cell is bursting regularly such as in control and in the 0.8*nA* condition, the interspike distributions consist of one large value that corresponds to the interburst interval (≈ 640msec in control) and several smaller values around 10msec which correspond to the ISI within a burst. There are current values for which the activity appears irregular and correspondingly, the ISI values are more diverse. Figure 5B shows the response of the model to larger depolarizing currents. The activity undergoes a sequence of interesting transitions which result in tonic spiking. When *I_e_* = 3.45*nA* the activity is periodic and there are 4 ISI values, larger currents result in 2 ISI values and tonic spiking produces one ISI value. Figure 5C shows the ISI distributions (y-axis, logarithmic scale) for each value of injected current (x-axis).

**FIG. 5.**
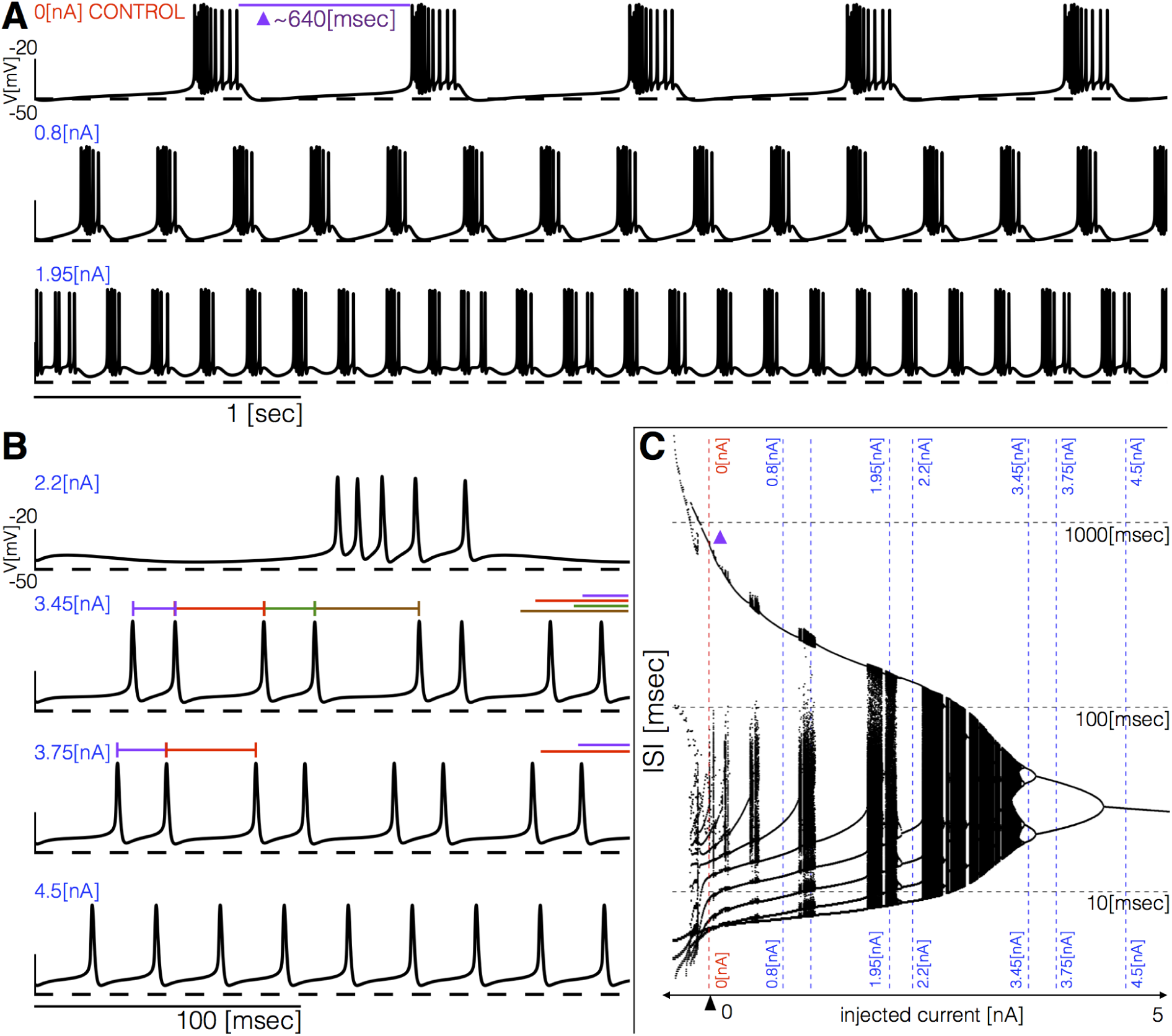
Response to current injections and interspike-intervals (ISI) distributions of model (a). **A** (top) Control traces (no current injected 0*nA*), regular bursting (0.8*nA*), irregular bursting 1.95*nA*. **B** (top) Fast regular bursting (*f_b_* ≈ *6Hz*), quadruplets (3.45*nA*), doublets (3.75*nA*) and singlets (4.5*nA*) (tonic spiking). **C** ISI distributions over a range of injected current.

All these bursters transition into tonic spiking regimes for depolarizing currents larger than *5nA* but they do so in different ways. To explore these transitions in detail, we computed the inter-spike interval (ISI) distributions over intervals of 60*sec* for different values of the injected current. Figure 6 shows the ISI distributions for the six models at *N* = 1001 equally spaced values of injected current over the shown range. The y-axis shows the values of all ISIs in logarithmic scale and the x-axis corresponds to injected current. In control, the ISI distribution consists of a few small values (< 100*msec*) that correspond to the ISIs of spikes within a burst, and a single larger value (> 100*msec*) that corresponds to the interval between the last spike of a burst and the first spike of the next burst. When the cell fires tonically the ISI distributions consist of a single value. The ISI distributions exhibit complicated dependences with the control parameter that result in beautiful patterns. For some current values, the cells produce small sets of ISI values indicating that the activity is periodic. However, this activity is quite different across regions. Interspersed with the regions of periodicity there are regions where the ISI distributions densely cover a band of values indicating non-periodic activity. Overall the patterns feature nested forking structures that are reminiscent of classical period doubling routes to chaos (Feigenbaum, 1978).

**FIG. 6.**
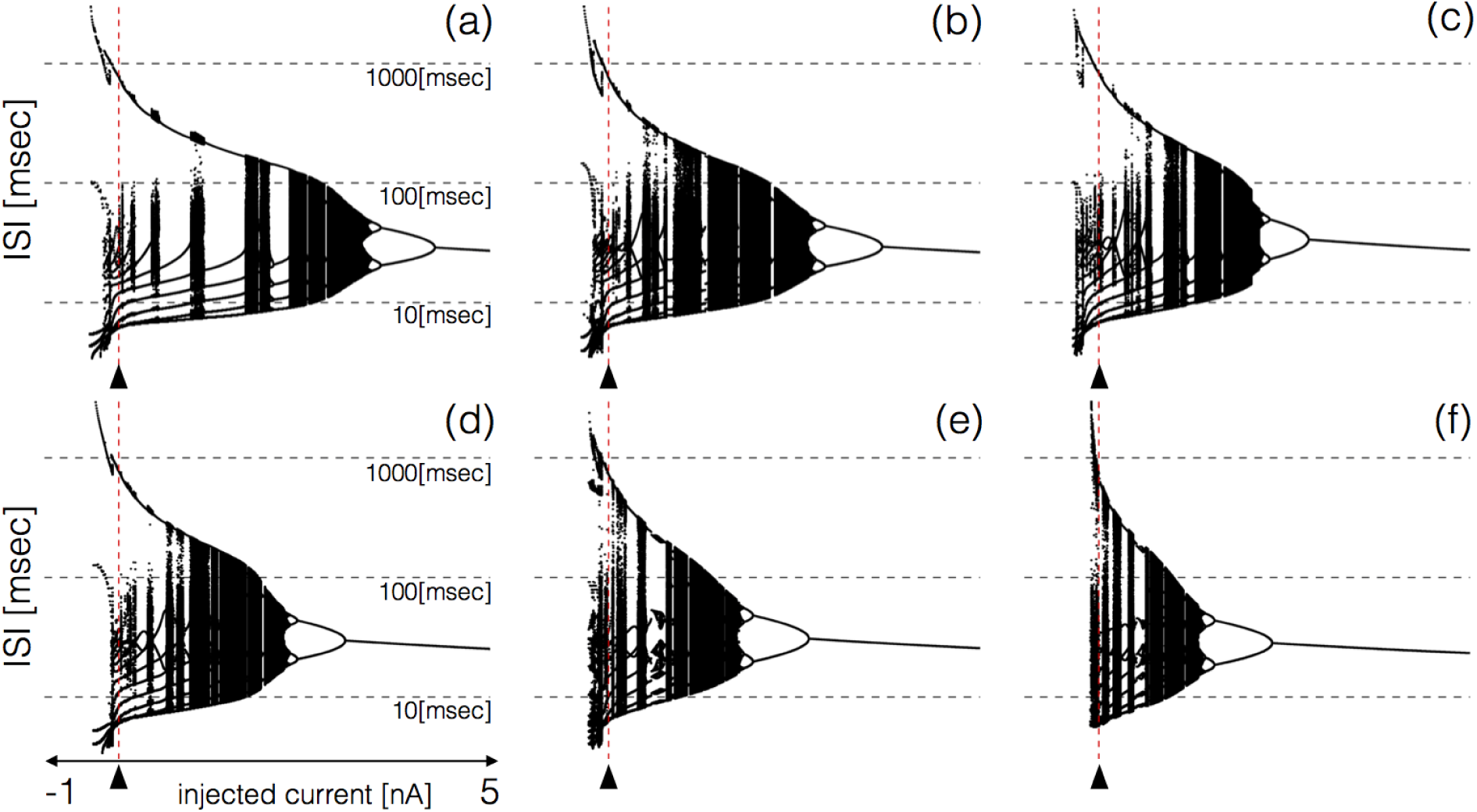
ISI distributions of the six model bursting neurons over a range of injected current. The panels show all ISI values of each model burster over a range on injected currents (vertical axis is logarithmic). All bursters transition into tonic spiking regimes for injected currents larger than *5nA* and the details of the transitions are different across models.

### Perturbing the models with gradual decrements of the maximal conductances

Figures 7 and 8 show the activity of the bursters when each of their channels are gradually decreased. We simulate the application of blockers by gradually decreasing the maximal conductances. The figures show 3 seconds of data for each condition. In all panels, the top traces correspond to the control condition (100%) and the traces below show the activity that results from decreasing the maximal conductance. The dashed lines are placed for reference at −50*mV* and 0*mV*. Each panel shows the traces for 11 values of the corresponding maximal conductance equally spaced between 100% (control) and 0% (completely removed). Each row of panels corresponds to a current type and the columns correspond to the different model bursters. Figure 7 displays the perturbations for the inward currents and Figure 8 shows the outward and leak currents.

**FIG. 7.**
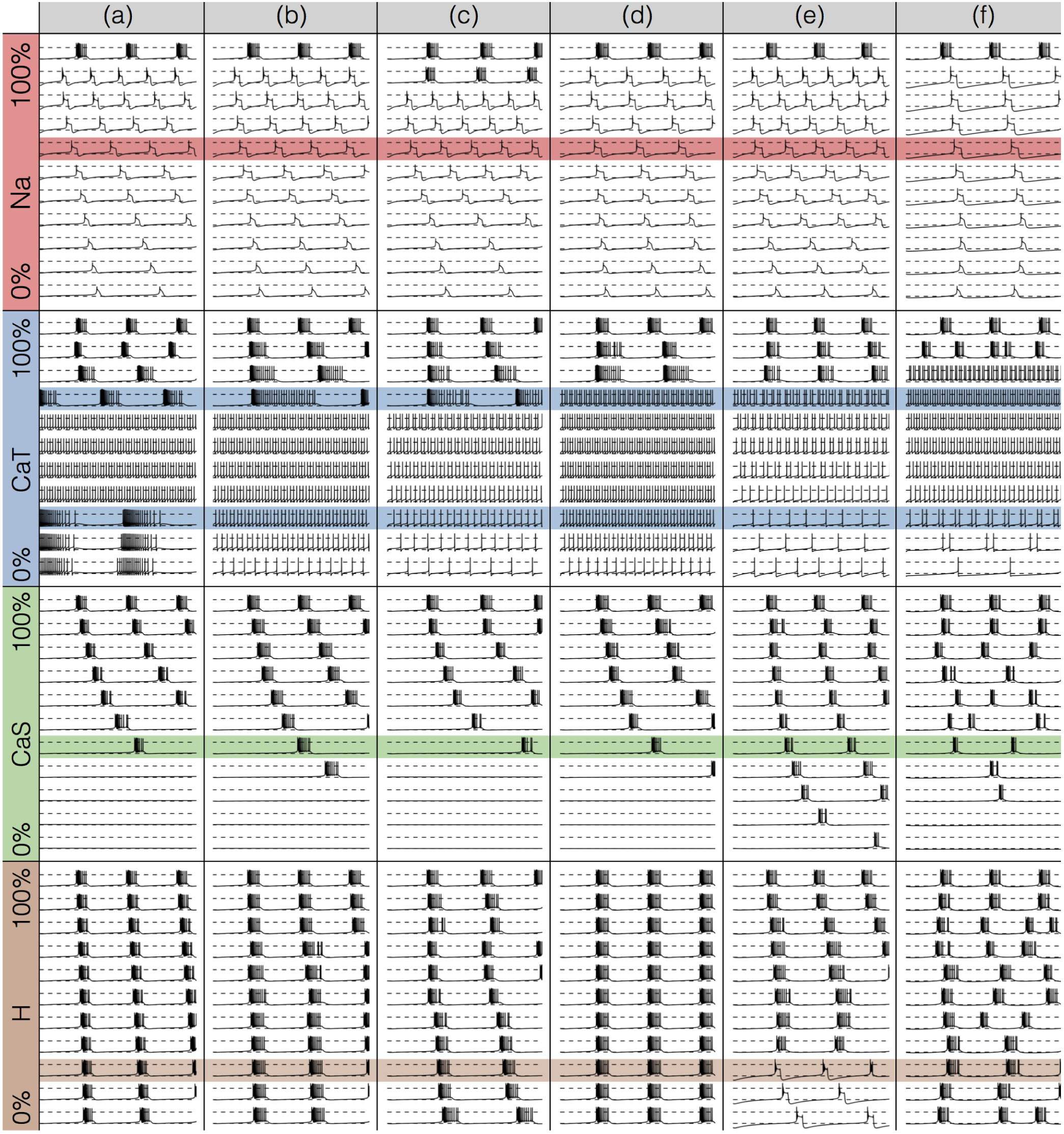
Effects of decreasing maximal conductances: inward currents. The figure shows the membrane potential *V* of all model cells as the maximal conductance *g_i_* of each current is gradually decreased from 100% to 0%. Each panel shows 11 traces with a duration of 3*secs*. Dashed lines are placed at 0*mV* and − 50*mV*. The shading indicates values of maximal conductance for which the activity the models differs the most.

**FIG. 8.**
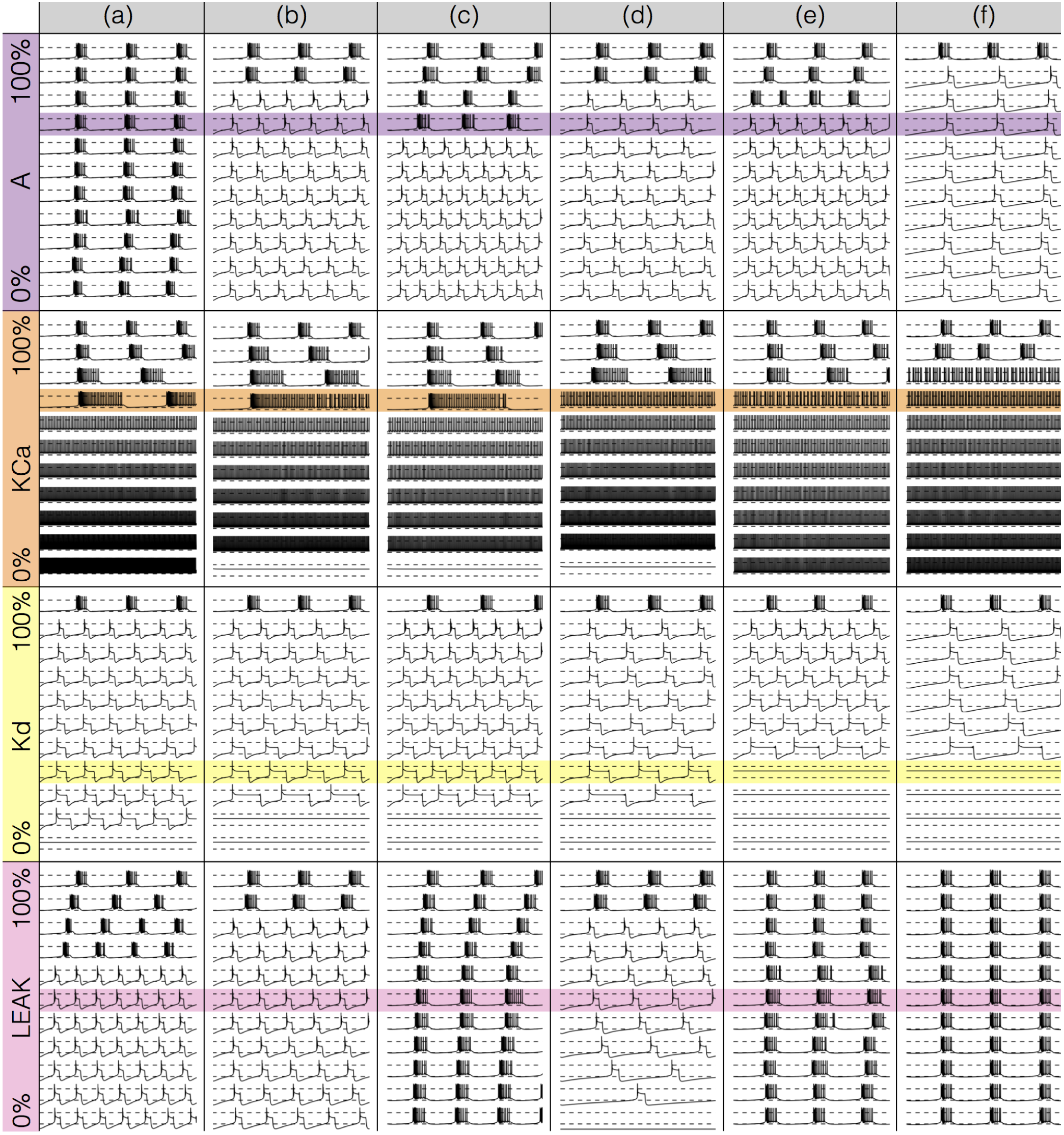
Effects of decreasing maximal conductances: outward currents. The figure shows the membrane potential *V* of all model cells as the maximal conductance *g_i_* of each current is gradually decreased from 100% to 0%. Each panel shows 11 traces with a duration of 3*secs*. Dashed lines are placed at 0*mV* and −50*mV*. The shading indicates values of maximal conductance for which the activity the models differs the most.

The top row in Figure 7 shows the effect of gradually decreasing the *Na* current in the bursters. In all cases, the activity transitions from the control bursting behavior to a single-spike bursting regime. These transitions occur at different levels and the final traces, when the *Na* current is completely removed (panel bottom), have different frequencies. The effect of decreasing *CaT* is less consistent across models. When the *CaT* is completely removed, most models transition into a tonic spiking regime but the spread of frequencies is much larger than in the *Na* case. The exception is model (a), which remains a burster but at a much lower frequency and with duty cycle ≈ 0.5. The intermediate decrements highlighted in blue also show visible differences. When *gCaT → 0.7gCaT* models (a), (b) and (c), show bursting activity at different frequencies and with different duty cycles. Models (d), (e) and (f), become tonic spikers at this condition, but their frequencies are different. Note that in the case of model (e) the spiking activity is not regular and the ISIs take several different values. When *gCaT → 0.2gCaT* most models spike tonically but now (e) is regular and (f) shows doublets. Model (a) is the exception and remains a burster.

Gradually removing *CaS* has a strong effect in the frequency of bursts and in most models this perturbation does not affect the burst duration or stability of the oscillations. This perturbation is most disruptive in model (f) where *CaS* dominates. When *gCaS → 0.4gCaS* all models remain bursting but they do so at different frequencies and the burst duration in (f) is halved. Gradually removing the *H* current disrupts the stability of the oscillations. Most models display bursts but the activity is no longer periodic. When *gH →* 0.2*gH* most models burst and their frequencies are similar to the controls. Model (e) is the exception and becomes a single-spike burster.

Figure 8 shows the effect of gradually decreasing the outward and leak currents. Decreasing the *A* channel results again in divergent responses across the population of bursters. Burster (a) is the only model that stays relatively unaffected by this perturbation, probably because *gA* is lowest in this model. The rest of the models transition into single-spike bursting regimes. This transition occurs at different simulated blocker concentrations and the frequencies of the blocked states are different. When *gA* → 0.7*gA* models (a) and (c) remain bursting while the rest depart visibly from their controls.

Decreasing *KCa* has a similar effect across models: burst frequencies decrease and burst durations increase up to a critical value after which the models spike tonically (with different frequencies). Decreasing *Kd* disrupts the models in a similar way: all models lose the capacity to produce bursts. The activity corresponds to single-spike bursting modes up to a critical decrease after which the models become quiescent. This transition occurs at different points across models. When *gKd* → 0.3*gKd* models (a), (b), (c) and (d) produce oscillations at different frequencies and models (e) and (f) are quiescent. Finally, decreasing the leak current leads to different behaviors across models. Models (c) and (e) remain relatively unaffected except for small changes in burst frequency (model (f) has no leak channel). In contrast, models (a), (b) and (d), transition into a single-spike bursting modes with different frequencies. This transition occurs at different conductance values. When the leak channel is completely removed, only model (d) becomes quiescent. Note that the membrane potential of the quiescent state is below −50*mV* unlike the *Kd* case where quiescence corresponds with more depolarized states.

### Effects of gradually removing *CaT* on all currents

Gradually removing one current impacts the dynamics of all currents. We illustrate this using currentscapes to inspect how the contributions of currents change in each condition. Figure 9 shows the currentscapes of model (f) as the maximal conductance of the *CaT* current is gradually decreased. Each panel corresponds to a different simulated blocker concentration and shows the membrane potential on top, and the currentscapes at the bottom. The top panels show 1 second of data and correspond to the 100%*gCaT* (control), 90%*gCaT* and 80%*gCaT* conditions. The center panels show 0.1 seconds of data for decrements ranging from 70% to 20% and the bottom panels show 2 seconds for the 10% and 0% conditions. As *CaT* is gradually removed the activity transitions from a bursting regime to a tonic spiking regime.

**FIG. 9.**
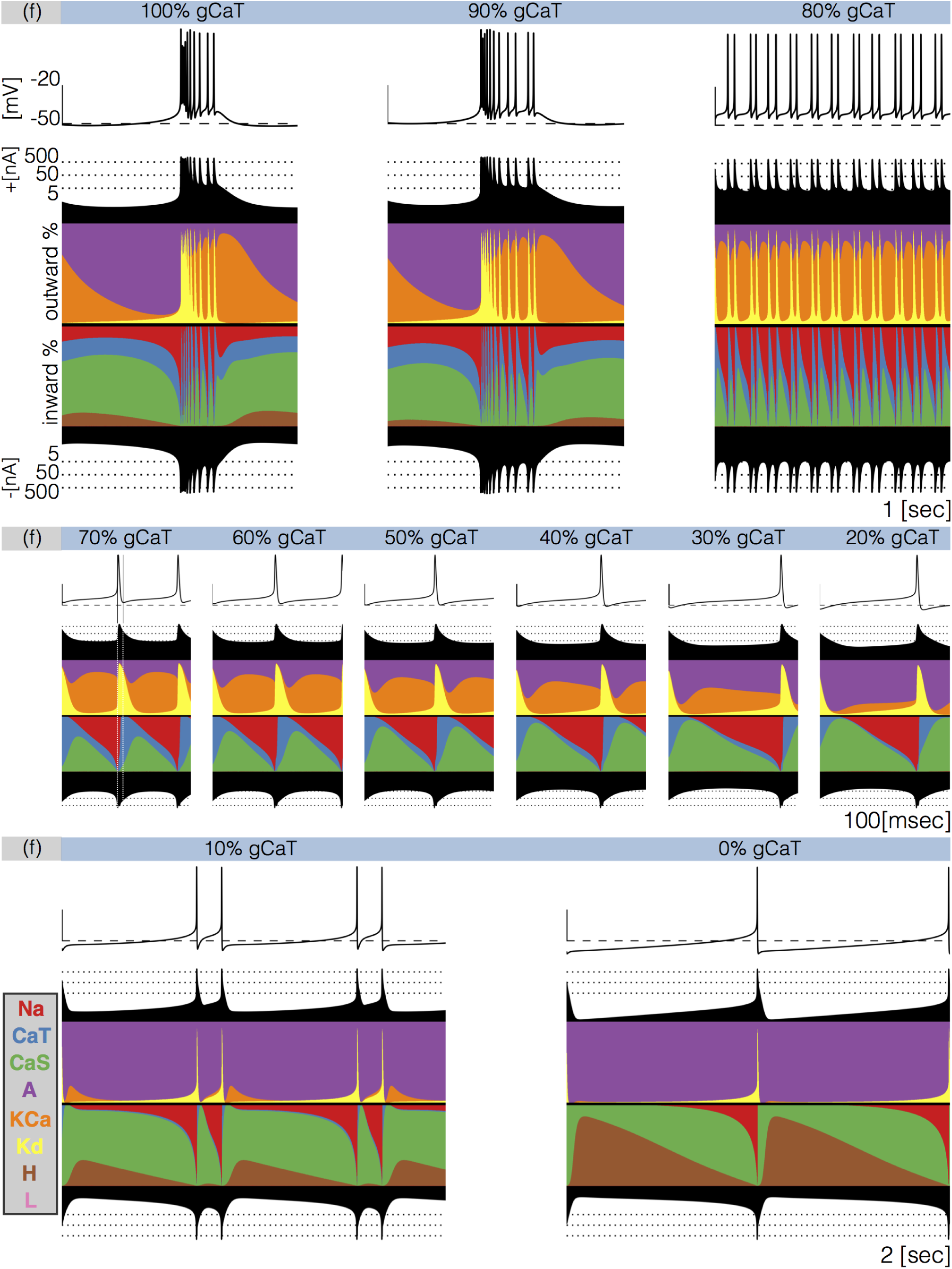
Decreasing *CaT* in model (f). The figure shows the traces and the currentscapes of model (f) as *CaT* is gradually decreased. Top panels show 1 second of data, center panels show 0.1 seconds and the bottom panels show 2 seconds (see full traces in Figure 8).

When *gCaT* → 90% the neuron produces bursts but these become irregular and their durations change. Decreasing the conductance to 80%*gCaT* results in completely different activity. The spiking pattern appears to be periodic but there are at least three different ISI values. It is hard to see changes in the *CaT* contribution across these conditions, but changes in other currents are more discernible. The contribution of the *A* current that is large in the control and 90%*gCaT* conditions and is much smaller in the 80%*gCaT* condition. Additionally, the *Na* and *KCa* currents show larger contributions, the *CaS* current contributes less and the *H* current is negligible. Further increments in simulated blocker concentration result in tonic spiking regimes with frequencies ranging from ≈ 20*Hz* to ≈ 10*Hz*. The center panels in Figure 9 show the currentscapes for these conditions on a different time scale to highlight the contributions of *CaT*. The leftmost panel shows the 70%*gCaT* condition. In this panel, we placed vertical lines indicating the time stamps at which the peak of the spike and the minimum occur. Notice the large contribution of the *Na* current prior to the peak of the spike, and the large contribution of the *Kd* current for the next ≈ 10*msec*. When the membrane potential is at its minimum value the *CaT* current dominates the inward currents and remains the largest contributor for the next ≈ 10*msec*. The *CaT* current reduces it share drastically by the time the *Na* current is visible and *CaS* takes over. The contribution of *CaT* remains approximately constant during repolarization and vanishes as the membrane becomes depolarized and the *Na* current becomes dominant. The effect of removing *CaT* is visible in this scale. The waveform of the contribution remains qualitatively the same: largest at the minimum voltage and approximately constant until the next spike. However, the contribution of *CaT* during repolarization becomes smaller, and for larger conductance decrements results in a thiner band. Finally, the bottom panels show the cases 10%*gCaT* and 0%*gCaT* which correspond to a two-spike burster and a tonic spiker respectively. Note that even though the contribution of *CaT* is barely visible, complete removal of this current results in a very different pattern. The activity switched from bursting to spiking and the current composition is different; *KCa* disappeared in the 0% condition and the *A* current takes over. Notice also the larger contribution of the *H* current.

### Effects of completely removing a current

Figures 10 and 11 show the currentscapes for the completely removed condition for all models and all channel types. The figures consist of 4 × 6 panels where the models are arranged along columns and the current types along rows. Each panel shows 2 seconds of data. The inward currents are portrayed in Figure 10 and the outward and leak currents are shown in Figure 11. The top row in Figure 10 shows the activity and the currentscapes of the bursters when the *Na* current is completely removed. All bursters become single-spike bursters but the frequencies and the contributions of each current to the activity are different across models. The calcium currents are sufficient to initiate a burst and elicit a spike in all models. Completely removing *CaT* (second row) results in tonic spiking in most models and here again the currents are different across models. Note that the activity is similar in models (b) and (d) and so are their currents. This is not the case for models (c) and (e), however, which also display similar activity but have different contributions from each current. Removing the *CaS* current (third row) leads to quiescence in most models, except for model (e) that remains bursting at a low frequency (*τ* ≈ 3 secs.). The currents are similar for models (b), (c), (d) and (f), with larger *A* contributions than in (a), and *H* contributions not present in (e). Removing the *H* current does not disrupt bursting in most models but burst durations are affected. The exception is model (e) that transitions into a single-spike bursting regime. Notice the small oscillation before hyperpolarization.

**FIG. 10.**
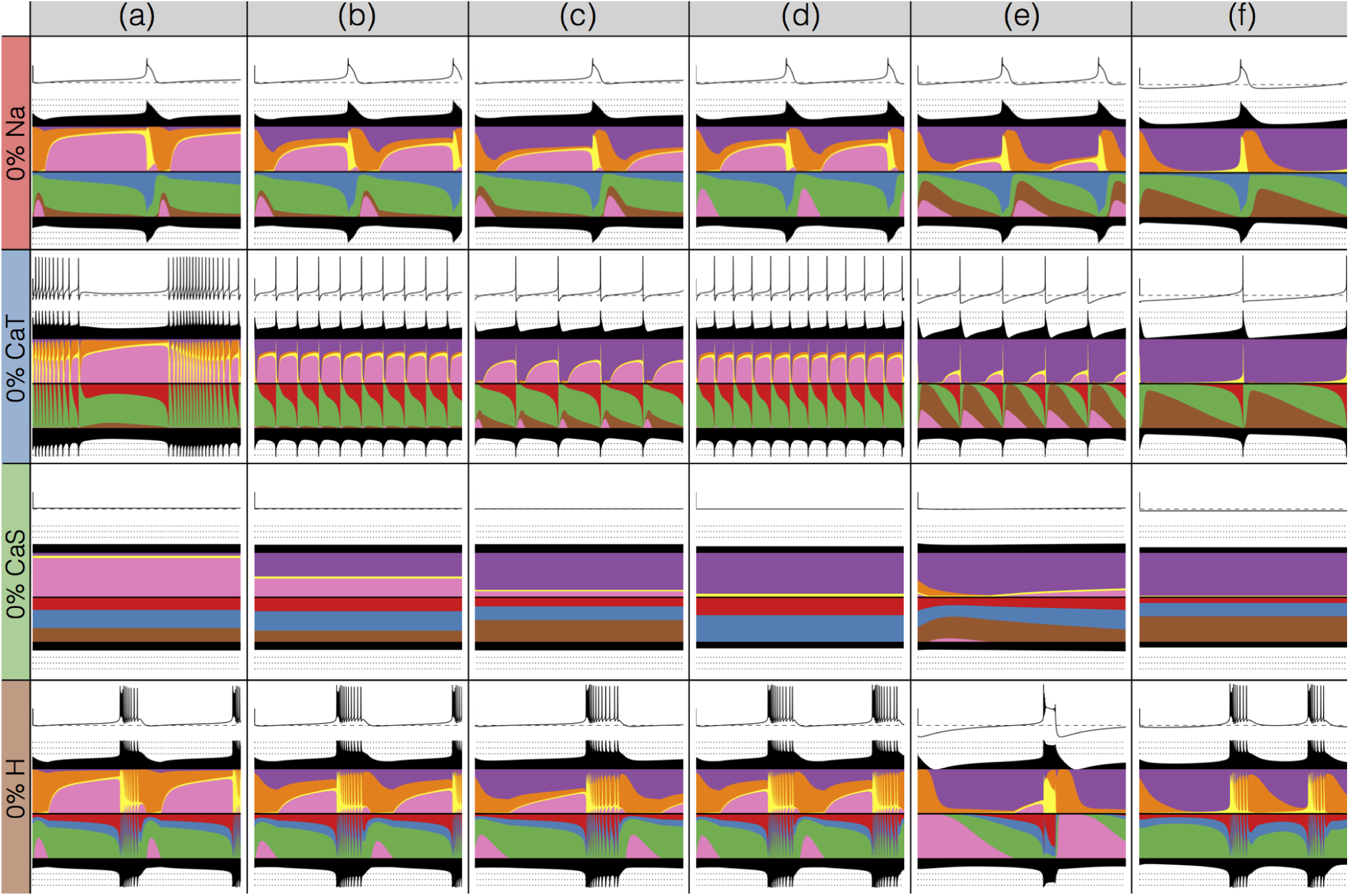
Complete removal of one current: inward currents. The figure shows the traces and currentscapes for all bursters when one current is completely removed.

**FIG. 11.**
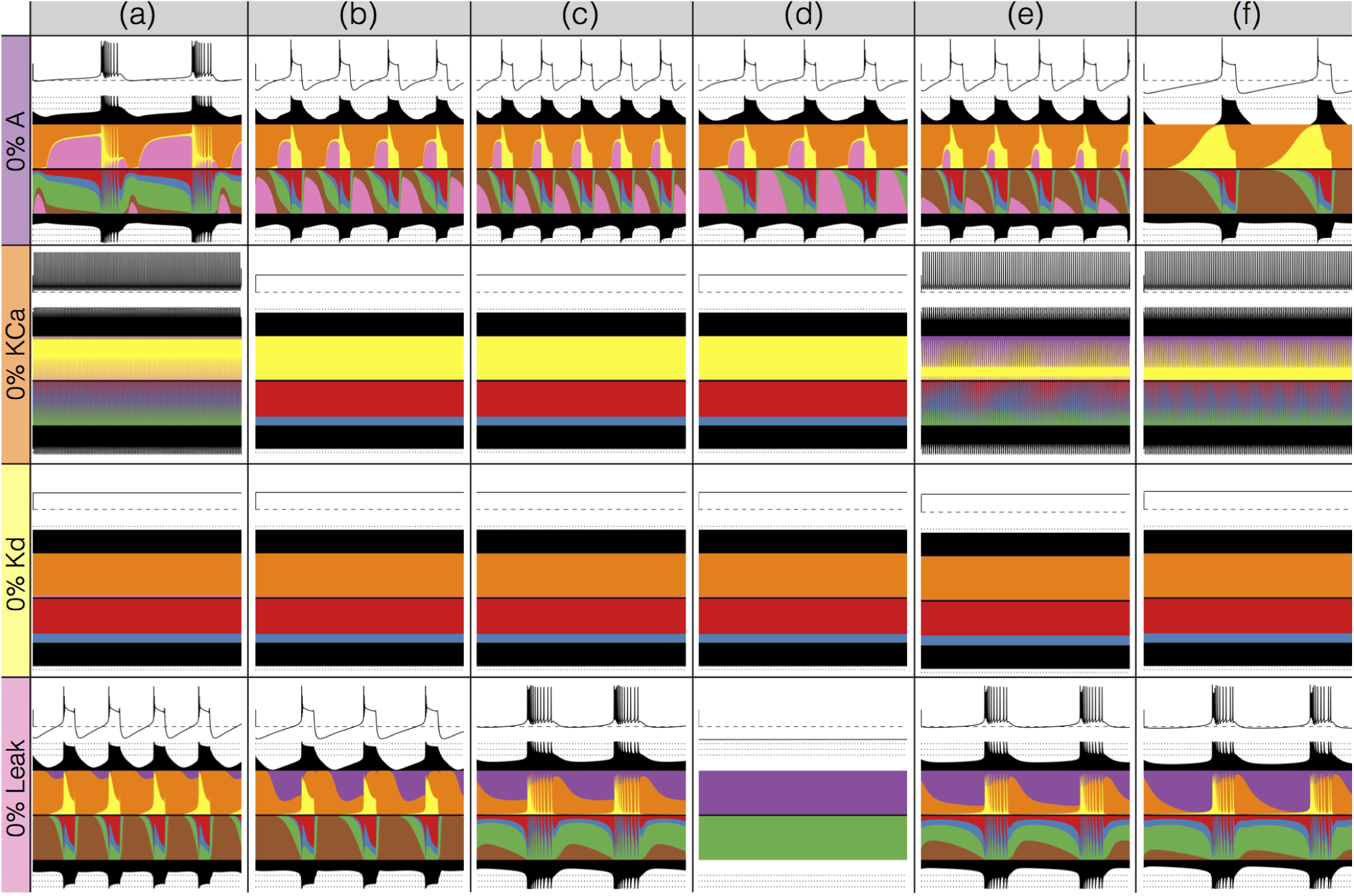
Complete removal of one current: outward currents. The figure shows the traces and currentscapes for all bursters when one current is completely removed.

The top row in Figure 11 shows the activity and the currentscapes of the bursters when the *A* current is completely removed. This perturbation results in similar activities across models. Model (a) has the smallest *gA* value and is virtually unaffected by this perturbation, except that the burst duration can be different from burst to burst. The rest of the models transition to single-spike bursting regimes with different frequencies. The activity in models (b), (c) and (d) is similar but the ratio of *H* and leak currents is different. Models (d) and (f), which do not have *leak* and *H* channels respectively, still display similar waveforms at a slower pace. Removing *KCa* results in tonic spiking in models (a), (e), and (f), and in quiescence in models (b), (c), and (d). In the case of the spikers the frequencies and the contributions of the *A* current are different. Removing the *Kd* current results in quiescence and in this case, the currents are distributed in almost identical proportions across models. Finally, removing the *leak* current result in different activities. Models (a) and (b) become single-spike bursters but their currents and frequencies are different. Bursters (c) and (e) increase their frequency but remain mostly unaltered despite the visible contributions of this current in the control conditions. Model (d) becomes quiescent and the total inward and outward currents show the lowest values (below < 1*nA*). Model (f) has no leak channel.

### Changes in waveform as conductances are gradually decreased

Figures 12 and 13 show how the waveform of the membrane potential *V* changes as currents are gradually decreased. The panels show the ridges of the probability distributions *p*(*V*) of the membrane potential *V*(*t*) for 1001 values of maximal conductance values (see methods). The probability of *V* (*t*) was computed using 30 seconds of data after dropping a transient period of 120 seconds. It was estimated using *N_b_* = 1001 bins in the range (−70, 35)*mv* and *N* ≈ 2 × 10^6^ samples for each maximal conductance value. The system spends more time in regions where 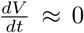 and is sampled more at those values. Therefore, features such as the amplitudes of the spikes appear as sharp peaks in the probability distributions. To highlight these peaks and visualize how they change as currents are gradually decreased, we plot the derivative or sharpness of the distribution in colors (see color scale in Fig. 3D).

**FIG. 12.**
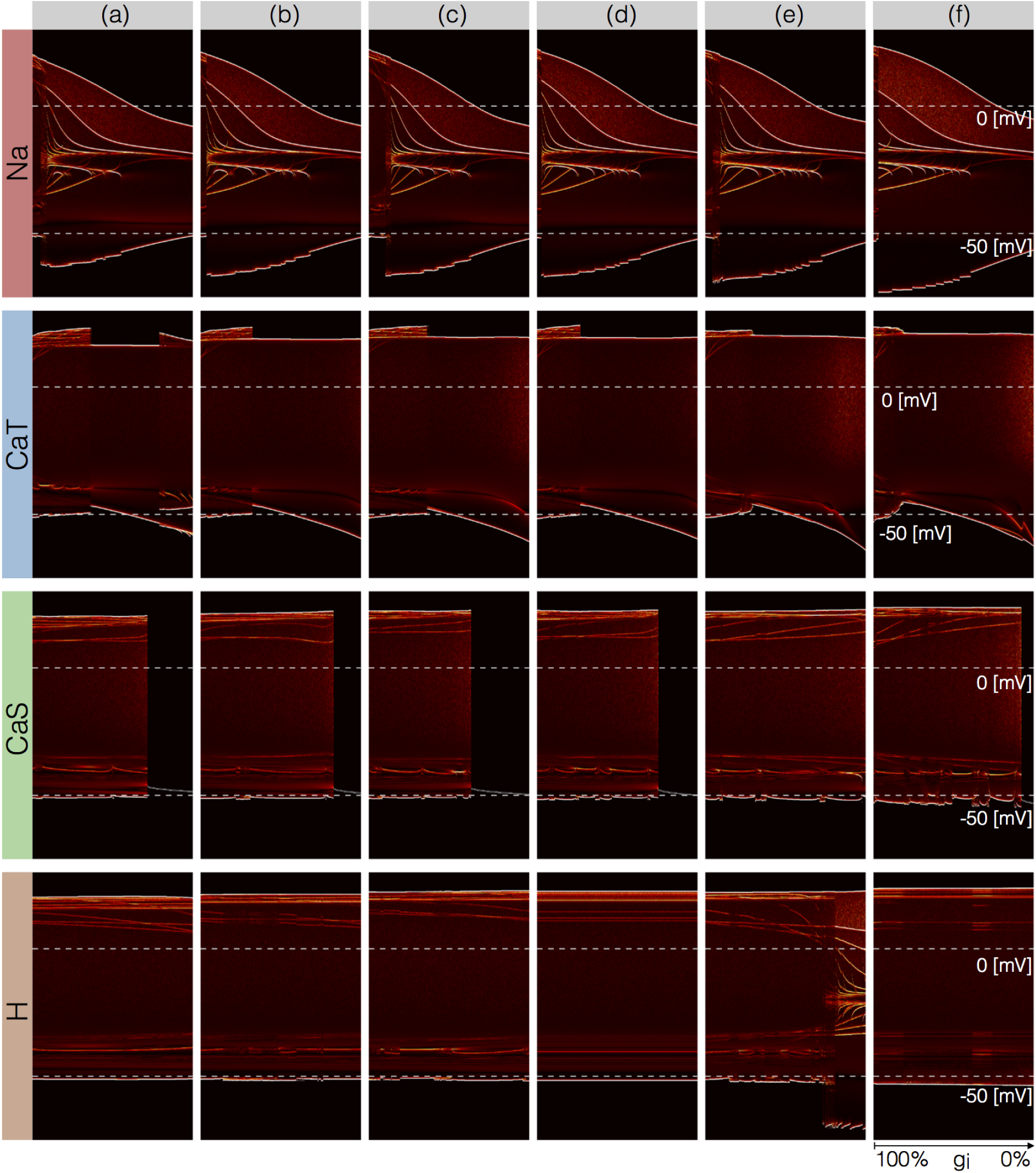
Changes in waveform as currents are gradually removed. Inward currents. The figure shows the ridges of the probability distribution of *V*(*t*) as a function of *V* and each maximal conductance *p*(*V, g_i_*). The ridges of the probability distributions appear as curves and correspond to values of V where the system spends more time, such as extrema. The panels show how different features of the waveform such as total amplitude, and the amplitude of each spike, change as each current is gradually decreased.

**FIG. 13.**
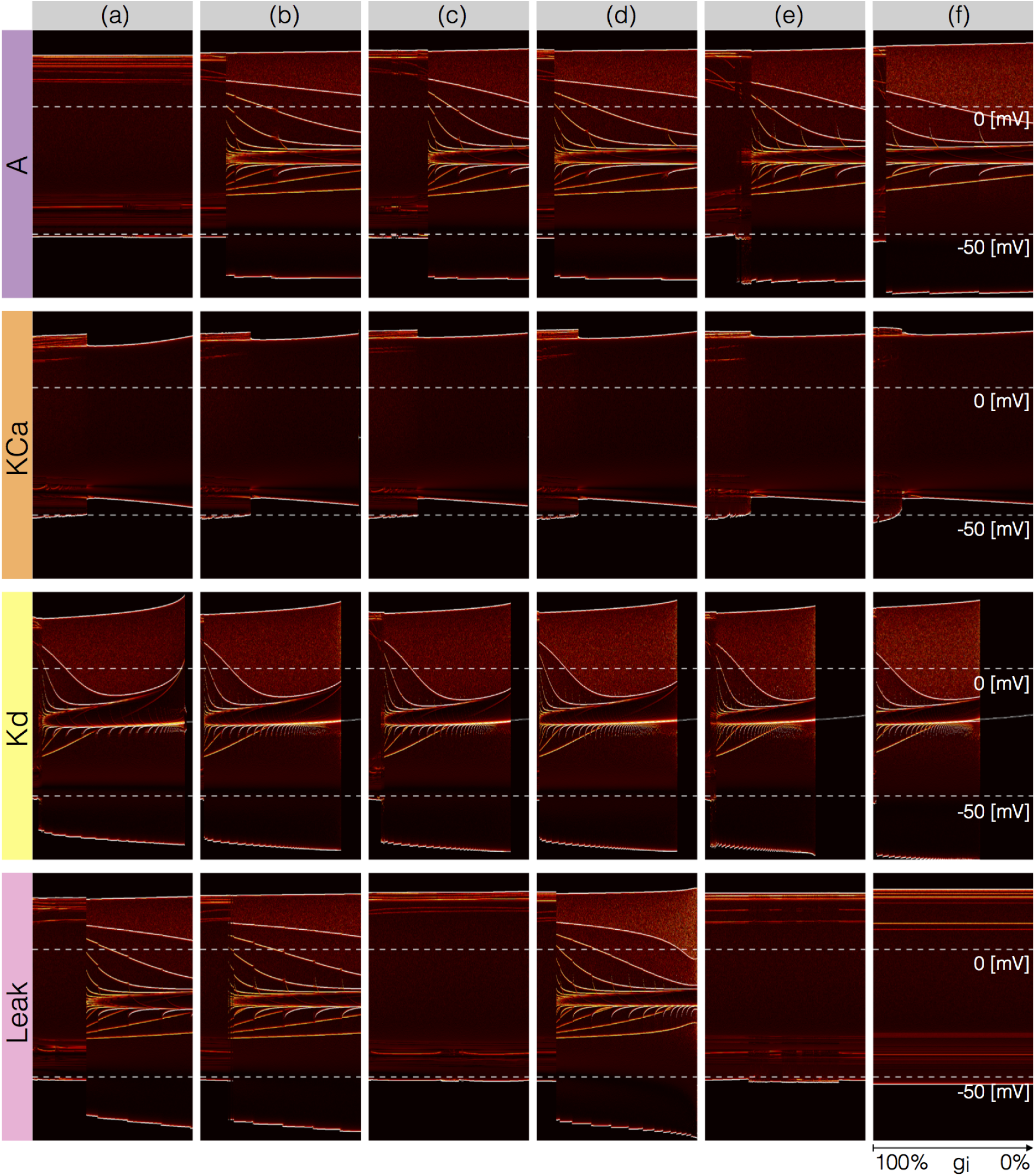
Changes in waveform as currents are gradually removed. Outward and leak currents. The figure shows the ridges of the probability distribution of *V*(*t*) as a function of *V* and each maximal conductance *p*(*V*; *g_i_*). See Figure 12.

Figure 12 shows this analysis for the inward currents. The top rows correspond to removing the *Na* current in the models. Note that the minimum value of *V* in control (left) is close to −50*mV* and a small increment in *gNa* results in a larger amplitude. The colored curves inside the envelopes correspond to the spikes’ amplitudes and features of the slow waves. For instance, when the *Na* current is completely removed (right) the amplitude of the oscillation is ≈ 40*mV* and the activity corresponds to a single-spike bursting mode. The spike amplitude is given by the edge of the colored region and the curve near *≈* − 20*mV* indicates the burst “belly”: the membrane hyperpolarizes slowly after spike termination and there is a wiggle at this transition. Removing *CaT* in model (a) does not disrupt bursting activity immediately. Notice that the amplitude of the bursts remains approximately constant over a range of concentrations. The dim red and yellow lines at ≈ 20*mV* show that the amplitudes of the spikes are different and have different dependences with *gCaT*. When the model transitions into a tonic spiking regime, the amplitude of the spikes is the same and there is only one amplitude value. This value stays constant over a range but the minimum membrane potential decreases and the overall amplitude therefore increases. The model returns to a bursting regime for values of *gCaT* smaller than 30%*gCaT*. Notice that in model (a) the membrane potential during bursts goes below −50*mV*, unlike in the control condition. Notice that the waveform of the membrane potential changes abruptly as *gCaT* is reduced and the models transition into a spiking regime. Model (f) is less resilient to this perturbation since this transition takes place at lower conductance values. Removing *CaS* does not much change the waveform, but it alters the temporal properties of the activity. The models remain bursting up to a critical concentration and the amplitude of the spikes do not change much. The features of the slow wave do not change much either except in model (f). Model (c) is less resilient to this perturbation since it becomes quiescent at lower concentrations than the other models. The effect of gradually removing *H* appears similar to *CaS* in this representation. In this case again, the morphology of the waveform is less altered than its temporal properties (except in model (e) where a transition takes place).

Figure 13 shows the same plots for the outward and leak currents. Removing the *A* current has little effect in model (a) and this translates into curves that appear as parallel lines indicating spikes with different amplitudes that remain unchanged. The rest of the models exhibit a transition into a different regime. The waveforms of this regime appears similar to the waveforms which result from removing *gNa* (see Fig. 7) but in this representation it is easier to observe differences such as the overall amplitude of the oscillation. The amplitude decreases as *gNa* is decreased and increases as *gA* is decreased. Removing *KCa* has a similar effect than removing *gCaT* in that the models transition into a tonic spiking regimes. The difference is that the spiking regimes which result from removing *KCa* have smaller amplitudes and also correspond to more depolarized states. In the case of the *KCa* current the model transitions into a tonic spiking regime that is different from the tonic spiking regime in the *gCaT* case. All models are very sensitive to removing *Kd* and a low values result in single-spike bursting modes with large amplitudes. Model (c) is most resilient to this perturbation and exhibits a large range with bursting modes. These oscillations break down in a similar way to the *Na* case and display similar patterns. As before, the top edge corresponds to the amplitude of the large spike and the curves in the colored region correspond to extrema of the oscillation. After spiking, the membrane remains at a constant depolarized value (≈ −20*mV*) for a long period and produces a high frequency oscillation before hyperpolarization. The amplitude of this oscillation increases as *Kd* is further decreased, and this results in a white curve that starts above 0*mV* and ends above 0*mV*. The beginning of this curve corresponds to a high frequency oscillation that occurs after spike termination. This type of activity is termed plateau oscillations and was reported in models of leech heart interneurons (Cymbalyuk and Calabrese, 2000) and in experiments in lamprey spinal neurons (Wang et al., 2014). These features are hardly visible in the traces in Fig. 8 and are highlighted by this representation. Finally, the Leak case appears as a mixture of the *Na* and *A* cases. The cells remain bursting over a range of values and some of them transition into a single-spike bursting mode that is different from the *KCa* case.

### Changes in current contributions as conductances are gradually decreased

The key to the visualization method in Figs. 12 and 13 is to consider *V*(*t*) not as a time series but as a stochastic variable with a probability distribution. The same procedure could be applied to the time series of each current. However, because the contributions of the currents are different at different times and at different maximal conductance values, it is not possible to display this information using the same scale for all channels. To overcome this we proceed as in the currentscapes and instead focus on the normalized currents or shares to the total inward and outward currents (the rows of matrices *Ĉ^+^* and *Ĉ*^−^, see methods). The current shares *Ĉ_i_*(*t*) correspond to the width of the color bands in the currentscapes and can also be represented by a time series that is normalized to the interval [0, 1]. The probability distribution of *Ĉ_i_*(*t*) permits displaying *changes* in the contributions of each current to the activity as one current is gradually removed. Interpreting these distributions is straightforward as before: the number of times the system is sampled in a given current share configuration is proportional to the time the system spends there.

Figure 14 shows the probability distributions of the current shares as *CaT* is gradually decreased in model (f) (see also Figure 9). The panels show the share of each current as *CaT* is gradually decreased and the probability is indicated in colors. In control the *Na* and *CaT* current shares are distributed in a similar way. Both currents can at times be responsible for ≈ 90% of the inward current, but most of the time they contribute with ≈ 20%. The *Na* current is larger right before spike repolarization and the *CaT* contributes with about ≈ 90% of the small (≈ 5nA) total inward current. For larger decrements the system transitions into tonic spiking and the contribution of the *Na* current is more evenly distributed over a wider range. The contribution of the *CaT* current is predominantly ≈ 15% and trends to zero as *gCaT* → 0. Note also that as the contribution of *CaT* decreases, the contribution of *CaS* increases to values larger than 75% while in control it contributes with ≈ 50%. The contribution of the *H* current is evenly distributed between 0% and ≈ 80%, it becomes negligible between ≈ 80% and ≈ 20% and becomes dominant after 20%. The *A* current behaves similarly to the *H*. It contributes with ≈ 90% of the (small ≈ 2*nA*) total outward currents right before burst initiation and its contribution decreases drastically when the system transitions into tonic spiking. As *CaT* is removed further the *A* current is more likely to contribute with a larger share. The contribution of the *KCa* current decreases as *gCaT* is decreased and some of it persists even when *gCaT* is completely removed. Finally, the contribution of the *Kd* current does not appear to change much and nor does its role in the activity.

**FIG. 14.**
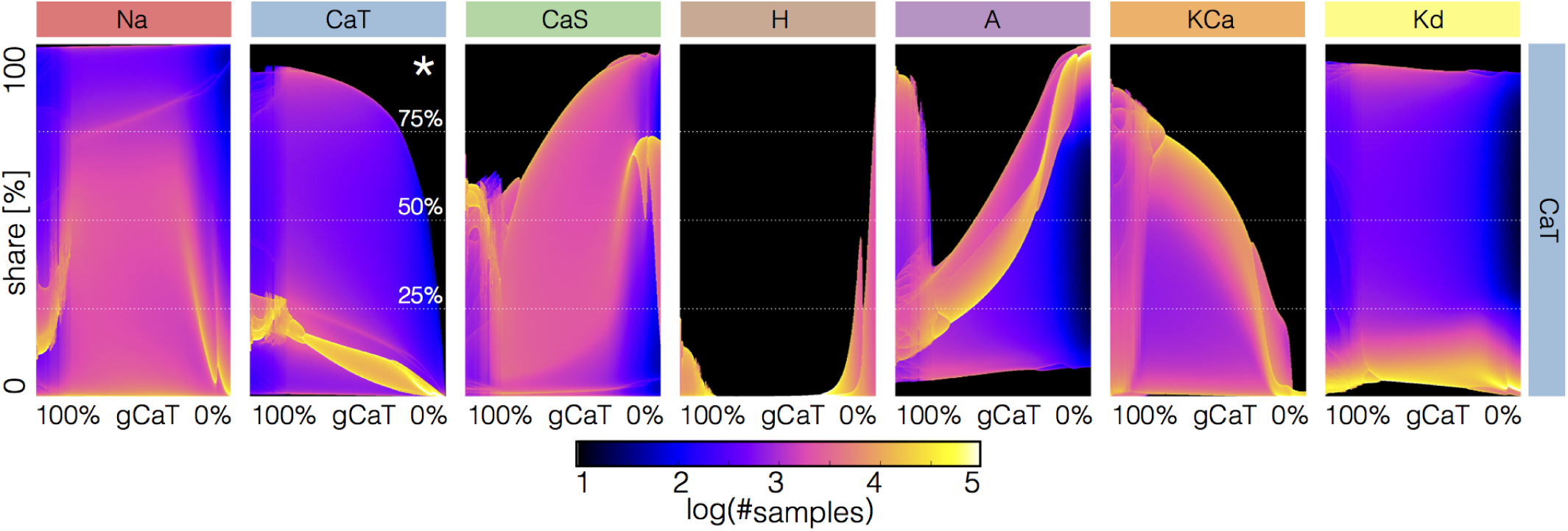
Changes in waveform of current shares as one current is gradually decreased. The panels show the probability distribution of the share of each current 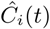 for model (f) as *CaT* is decreased

Figure 15 shows the effect of gradually decreasing each current on all currents in model (c). The figure features 8 × 8 = 64 panels. The rows indicate which channel is removed and the columns show the effect of this perturbation on all the currents. The first row shows how the shares of each current to the activity change as the *Na* current is decreased. For instance, the effect of decreasing *gNa* on the *Na* current (indicated by *) is as expected, with the maxima of the distribution trending to zero as *gNa* → 0. The effect of removing *gNa* on the other currents is non-trivial and is displayed along the same row. The rows show the same analysis for other channels indicated in the figure label. Notice that while the effect of removing a current on that same current (diagonal panels) is relatively predictable, the rest of the currents become rearranged in complicated ways. Finally, note that in all cases for small decrements there are no negligible currents since for each of them, there are periods of time where they contribute to at least ≈ 20% of the total current.

**FIG. 15.**
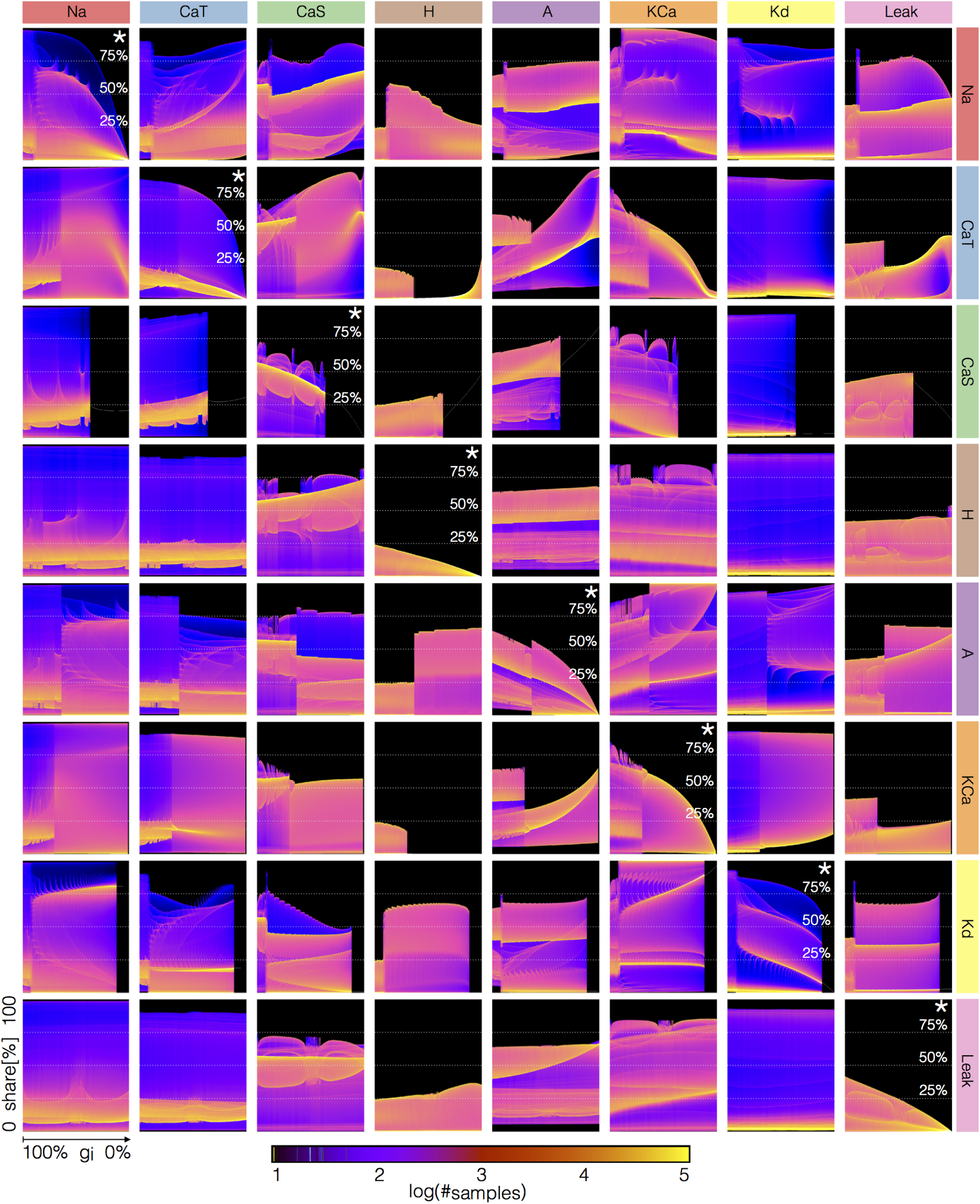
Changes in waveform of current shares as one current is gradually decreased. The panels show the probability distribution of the share of each current *Ĉ_i_*(*t*) for model (c) as each current is decreased

## III. Discussion

There is an ever larger availability of experimental data to inform detailed models of identified neuron types (Mc-Dougal et al., 2017). Experimenters have determined the kinetics of many channel types both in vertebrate and invertebrate neurons. There are also model databases with thousands of parameters which permit the development of large scale models of neural tissue (Bezaire et al., 2016). One difficulty in ensemble modeling is the necessity of incorporating the biological variability observed in some of the parameters – such as the conductances – at the same time that we require the models to capture some target activity. In other words, we may be interested in modeling a type of cell that displays some sterotypical behavior, and would like to obtain as many different versions of such models. Two main avenues to this problem were introduced in the past. One consists of building a database of model solutions over a search domain and screening for target solutions: this considers all possible value combinations within an allowed range up to a numerical resolution and then applies quantitative criteria to determine which solutions correspond to the target activity (Prinz et al., 2004). An alternative approach consists of designing a target function that assigns a score to the models' solutions in such a way that lower scores correspond to solutions that meet the targets, and then optimizing these functions (Achard and De Schutter, 2006; Druckmann et al., 2007; Ben-Shalom et al., 2012).

Both approaches have advantages and shortcomings. In the case of the database approach, trying all posible parameter combinations in a search range becomes prohibitively expensive as more parameters are allowed to vary. One advantage of this approach is that it provides a notion of how likely it is to find conductances within a search range that will produce the activity. In the landscape approach we find solutions by optimization and – without further analysis – we do not know how likely is a given solution type. This approach has the advantage that it can be scaled to include large numbers of parameters. Additionally, if a particular solution is interesting, we can use genetic algorithms on successful target functions to “breed” as many closely related models as desired. Ultimately, any optimization heuristic requires blind testing random combinations of the parameters, and developing quantitative criteria for screening solutions in a database results in some sort of score function, so both approaches are complementary. A successful target function can determine if a random perturbation results in disruption of the activity and this can be used to perform population-based sensitivity analyses (Devenyi and Sobie, 2016).

Regardless of the optimization approach most work is devoted to the design of successful target functions. Different modeling problems require different target functions (Roemschied et al., 2014; Fox et al., 2017; Migliore et al., 2018) and one challenge in their design is that sometimes we do not know a priori if the model contains solutions that will produce good minima. In addition, a poorly constrained target function can feature multiple local minima that could make the optimization harder, so even if there are good minima they may be hard to find. One difference between the landscape functions in Achard and De Schutter (2006) and the ones utilized here is the way that model solutions are compared to a target *time series*. They utilize a phase-plane method to compare the waveform of the membrane potential to some target time series. The functions introduced here use an analysis based on Poincaré sections or thresholds to characterize the waveform and to define an error or score. Instead of targeting a particular waveform we ask that some features of the waveform – such as the frequency and the burst duration – are tightly constrained, while other features – such as the concavity of the slow waves – can be diverse. This is motivated by the fact that across individuals and species, the activity of the pyloric neurons can be diverse but the neurons always fire in the same sequence and the burst durations have a well-defined mean. Our approach is successful in finding hundreds of models that display a target activity in minutes using a commercially available desktop computer. Application of evolutionary techniques to optimize these functions provides a natural mean to model the intrinsic variability observed in biological populations.

One of the main benefits of computational modeling is that once a behavior of interest is successfully captured we then possess a mechanistic description of the phenomena that can be used to test ideas and inform experiments (Coggan et al., 2011; Lee et al., 2016; Devenyi and Sobie, 2016; Gong and Sobie, 2018). As the models gain biophysical detail these advantages wane in the face of the complexity imposed by larger numbers of variables and parameters. Numerical models can generate large ammounts of data that can be hard to visualize and interpret. The development of novel visualization procedures has the potential to greatly assist intuition into the details of how these models work (Gutierrez et al., 2013). Here we introduced a novel representation of the dynamics of the ionic currents in a single compartment neuron. Our representation is simple and displays in a concise way the contribution of each current to the activity. This representation is easily generalizable to multi-compartment models and small networks.

We employed these procedures to build many similar bursting models with different conductance densities and to study their response to perturbations. The responses of the models to current injections and gradual decrements of their conductances can be diverse and complex. Inspection of the ISI distributions revealed wide ranges of parameter values for which the activity appears irregular, and similar regimes can be attained by gradually removing some of the currents. Period doubling routes to chaos in neurons have been observed experimentally and in conductance-based models (Hayashi et al., 1982; Hayashi and Ishizuka, 1992; Szucs et al., 2001; Canavier et al., 1990). The sort of bifurcation diagrams displayed by these models upon current injection are qualitatively similar to those exhibited by simplified models of spiking neurons for which further theoretical insight is possible (Touboul and Brette, 2008). Period doubling bifurcations and low dimensional chaos arise repeatedly in neural models of different natures including rate models(Ermentrout, 1984; Alonso, 2017). The bursters studied here are close (in parameter space) to aperiodic or irregular regimes suggesting that such regimes are ubiquitous and not special cases.

Overall our results show that in these model neurons, similar membrane activities can be attained by multiple mechanisms and that these underlying differences correspond to different current compositions at each stage of the cycle. Because the dynamical mechanisms driving the activity are different in different models, perturbations can result in qualitatively different scenarios. Our visualization methods allow us to gather intuition on how different these responses can be and to explore the contribution of each current type to the neural activity. Even in the case of single compartment bursters, the response to perturbations of a population can be diverse and hard to describe. To gain intuition into the kind of behaviors the models display upon perturbation, we developed a representation based on the classical probability of the membrane potential *V*. This representation permits displaying changes in the waveform of *V* as each current is blocked. This representation successfully captures the important observation that the models respond to simulated blockers in different ways, but that there are also similarities among their responses. A concise representation of the effect of a perturbation is a necessary step towards developing a classification scheme for the responses.

## IV. Methods

### Model equations

The membrane potential *V* of a cell containing *N* channels and membrane capacitance *C* is given by:

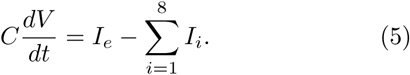

Each term in the sum corresponds to a current *I_i_* = *g_i_m^pi^h^qi^*(*V* − *E_i_*) and *I_e_* is externally applied current. The maximal conductance of each channel is given by *g_i_*, *m* and *h* are the activation and inactivation variables, the integers *p_i_* and *q_i_* are the number of gates in each channel, and *E_i_* is the reversal potential of the ion associated with the i-th current. The reversal potential of the Na, K, H and leak currents were kept fixed at *E_Na_* = 30*mV*, *E_K_* = −80*mV*, *E_H_* = −20*mV* and *E_leak_* = −50*mV* while the calcium reversal potential *E_Ca_* was computed dynamically using the Nernst equation assuming an extracellular calcium concentration of 3 × 10^3^*μM*. The kinetic equations describing the seven voltage-gated conductances were modeled as in Liu et al. (1998),

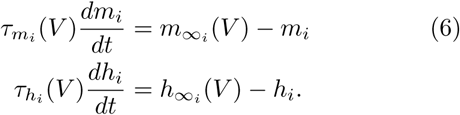

The functions 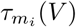, 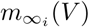, 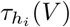 and 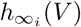 are based on the experimental work of Turrigiano et al. (1995) and are listed in refs. (Liu et al., 1998; Turrigiano et al., 1995). The activation functions of the *K_Ca_* current require a measure of the internal calcium concentration [*Ca*^+2^] (Liu et al., 1998). This is an important state variable of the cell and its dynamics is given by,

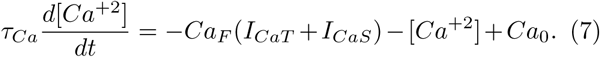

Here 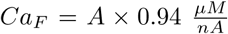 is a current-to-concentration factor and *Ca*_0_ = 0.05 *μM* (A = 0.628). These values were originally taken from Liu et al. and were kept fixed. Finally, *C* = 10*nF*. The number of state variables or dimension of the model is 13. We explored the solutions of this model in a range of values of the maximal conductances and calcium buffering time scales. The units for voltage are *mV*, the conductances are expressed in *μS* and currents in *nA*. Voltage traces were obtained by numerical integration of eq. (5) using a Runge-Kutta order 4 (RK4) method with a time step of *dt* = 0.1*msec* (Press et al., 1988).

### Currentscapes

The currentscapes are stacked area plots of the normalized currents. Although it is easy to describe their meaning, a precise mathematical definition of the images in Figure 2 can appear daring in a first glance. Fortunately, the implementation of this procedure results in simple python code.

The time series of the 8 currents can be represented by a matrix *C* with 8 rows and 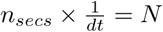 columns. For simplicity, we give a formal definition of the currentscapes for positive currents. The definition is identical for both current signs and is applied independently for each. We construct a matrix of positive currents *C*^+^ by setting all negative elements of *C* to zero, 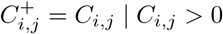 and 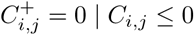. Summing *C*^+^ over rows results in a normalization vector *n*^+^ with *N* elements 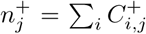. The normalized positive currents can be obtained as *Ĉ^+^* = *C^+^/n^+^* (element by element or entry-wise product). Matrix *Ĉ*^+^ is hard to visualize as it is. The columns of *Ĉ^+^* correspond to the shares of each positive current and can be displayed as pie charts. Here, instead of mapping the shares to a pie we map them to a segmented vertical “churro”. The currentscapes are generated by constructing a new matrix *C_S_* whose number of rows is given by a resolution factor *R* = 2000, and the same number of columns *N* as *C*. Each column *j* of *Ĉ^+^* produces one column *j* of *C_S_*. Introducing the auxiliary variable = 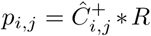 we can define the currentscape as,

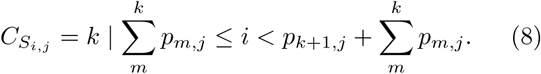

The integer *k ∈* [0, 7] indexes the current types and we assume 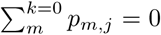. The black filled curve in Figure 2B corresponds to the normalization vector *n*^+^ plotted in logarithmic scale. We placed dotted lines at 5*nA*, 50*nA* and 500*nA* for reference throughout this work. The currentscapes for the negative currents are obtained by applying definition (8) to a matrix of negative *C^−^* currents defined in an analogous way as *C*^+^. Finally, note that matrices *Ĉ*^+^ and *Ĉ*^−^ are difficult to visualize as they are. The transformation given by definition (8) is useful to display their contents.

### ISI distributions

We inspected the effects of injecting currents in our models by computing the inter-spike interval ISI distributions. For this we started the models from the same initial condition and simulated them for 90 seconds. We dropped the first 30 seconds to remove transient activity and kept the last 60 seconds for analysis. Spikes were detected as described before. We collected ISI values for *N* = 1001 values of injected current equally spaced between − 1*nA* and 5*nA*.

### V distributions

To sample the distributions of *V* we simulated the system with high temporal resolution (*dt* = 0.001*msec*) for 30 seconds, after dropping the first 120 seconds to remove transients. We then sampled the numerical solution at random time stamps and kept 2 × 10^6^ samples *V* = {*V_i_*}.

**Author contributions**

LA: design of study, carried out simulations, prepared figures, wrote and edited the manuscript. EM: design of study, wrote and edited the manuscript.

**Acknowledgments** Research supported by MH046742 and R35 NS097343 (EM), T32 NS07292 and Swartz Foundation 2017 (LA).

## V. Acknowledgments

LMA acknowledges Marcos Trevisan for early discussions on genetic algorithms and Francisco Roslan for programming training.

